# Multi-areal neural dynamics encode human decision making

**DOI:** 10.1101/2022.08.05.502992

**Authors:** Jacqueline A. Overton, Karen Moxon, Matthew P. Stickle, Logan M. Peters, Jack J. Lin, Edward F. Chang, Robert T. Knight, Ming Hsu, Ignacio Saez

## Abstract

Value-based decision-making involves multiple cortical and subcortical brain areas, but the distributed nature of neurophysiological activity underlying economic choices in the human brain remains largely unexplored. Here, we use intracranial recordings from neurosurgical patients to show that risky choices are reflected in high frequency activity distributed across multiple prefrontal and subcortical brain regions, whereas reward-related computations are less widely represented and more modular. State space modeling reveals sub-second neural dynamics underlying choices during deliberation and allows high-accuracy trial-by-trial decoding of subjects’ choices robustly across patients despite differences in anatomical coverage. These results shed light into the neural basis of choice across brain areas and open the door to new intracranial approaches for brain state decoding.

## Introduction

Human decision-making involves the coordinated activity of multiple brain areas, acting in concert to adaptively generate choices. Abundant evidence implicates a variety of prefrontal and subcortical regions in this process, often conceptualized as a sequential transformation from abstract valuation to concrete motor plans(1–3). However, increasing evidence from animal models supports a distributed view of neural activity underlying reward-guided behavior, with activation across brain regions emerging simultaneously rather than sequentially during choice processes(4–7). In the human brain, little is known about how distributed neural activity enables adaptive choice. Moreover, the neurophysiological representation of choice-related signals across human brain areas remains mostly unknown, partially due to the difficulty of directly observing human neural activity with sufficient anatomical precision and temporal resolution. For example, whereas neural activity in multiple frequency bands, from slow oscillations to high-frequency activity are implicated in a variety of cognitive processes including memory(8–10), spatial navigation(11–13), attention(14, 15) and decision-making(16–18), a description of their relative contribution to generating choices, including their precise anatomical basis, is lacking. From a computational standpoint, earlier studies have shown that localized neural activity in a variety of regions reflects reward-related computations such as risk(19, 20) and reward probability(21, 22), but whether they co-occur with a distributed representation of choices is not well understood. Therefore, an anatomically accurate, temporally precise depiction of distributed neurophysiological activity during choice behavior is necessary.

Here, we leveraged human intracranial electroencephalography (iEEG) recordings from neurosurgical patients to study the relationship between distributed neurophysiological activity and choice behavior while overcoming some of the limitations of non-invasive human neural methodologies such as limited signal-to-noise ratio, temporal resolution (fMRI), or anatomical accuracy (EEG)(23, 24). Patients played an economic risky decision-making game during simultaneous multi-areal iEEG recording from multiple reward-related brain areas, including orbitofrontal, lateral prefrontal, premotor and motor cortices, insula, and amygdala. We examined iEEG local field potential (LFP) activity, which provides a direct readout of neural activity with high temporal resolution and anatomical precision, to examine the neurophysiological basis of choices across neural frequencies and regions and its temporal evolution prior to choice. Because decisions require representation of information related to both the utility of the presented choices as well as the sensorimotor aspects of the task, we also examine the regional representation of select reward computations, such as risk and win probability. In addition, we used linear dynamical systems models (LDS), capable of capturing the main neural dynamic features of distributed neural activity, to obtain further insights into the sub-second temporal dynamics and their relationship to behavior (i.e. choice evolution) and to build decoding models that allow decoding of overt behavior from neural dynamics alone.

Our results show that choice information (i.e., whether to choose a safe bet or a risky gamble) is represented primarily in higher-frequency activity (beta, gamma, high-frequency activity) across multiple cortical and subcortical brain areas. In contrast, individual choice-related computations (e.g., risk, win probability) are more unequally distributed across brain areas, with a more modular profile. LDS models capture the evolution of multi-areal brain activity during deliberation to reveal the sub-second evolution of choice dynamics and produce high trial-by-trial choice decoding accuracy from distributed iEEG activity in all patients in our sample (average = 74.6±3.19%), indicating that the combination of iEEG and LDS is robust to the high degree of variation in surgical strategy and anatomical electrode location.

Altogether, our results provide novel evidence for robustly decodable distributed high-frequency activity across multiple human brain areas during decision-making behavior, and a balance between distributed and localized brain processes underlying human choices. These results provide novel insights into the neurophysiological nature of human choices, and represent an important step to building general brain state decoders using intracranial approaches, which hold translational promise for disorders in which choice processes are altered, such as depression or addiction.

## Results

We recorded intracranial electroencephalography (iEEG) data from 36 medication refractory epilepsy patients while they played a gambling task(17), postoperatively during their stay at the epilepsy monitoring unit. Twenty patients had behavioral data of sufficient quality to be included in final analyses (see Methods). Patients made trial-by-trial choices between a safe prize and a risky gamble with varying win probabilities (see Fig. 1a). Patients gambled more often in trials with higher win probability, as expected in reward-maximizing behavior(25, 26) (Fig. 1b and Extended Data Fig. 1). Patients underwent either electrocorticography (ECoG), providing subdural coverage predominantly in frontoparietal regions (9/20 patients, Fig. 1c right), or stereotactic EEG (sEEG), predominantly in deep temporal lobe regions (amygdala, hippocampus, insula; 11/20 patients; Fig. 1c left and center, and Extended Data Fig. 2). We analyzed electrophysiological recordings from electrodes located in regions involved in reward-related behavior: orbitofrontal cortex (OFC), lateral prefrontal cortex (LPFC), cingulate cortex (CC), precentral gyrus (PrG), postcentral gyrus (PoG), parietal cortex (PC), amygdala (Amy), hippocampus (Hipp), and insula (Ins) (n=1085 electrodes total; Fig. 1c and Extended Data Table 1).

**Figure 1.**
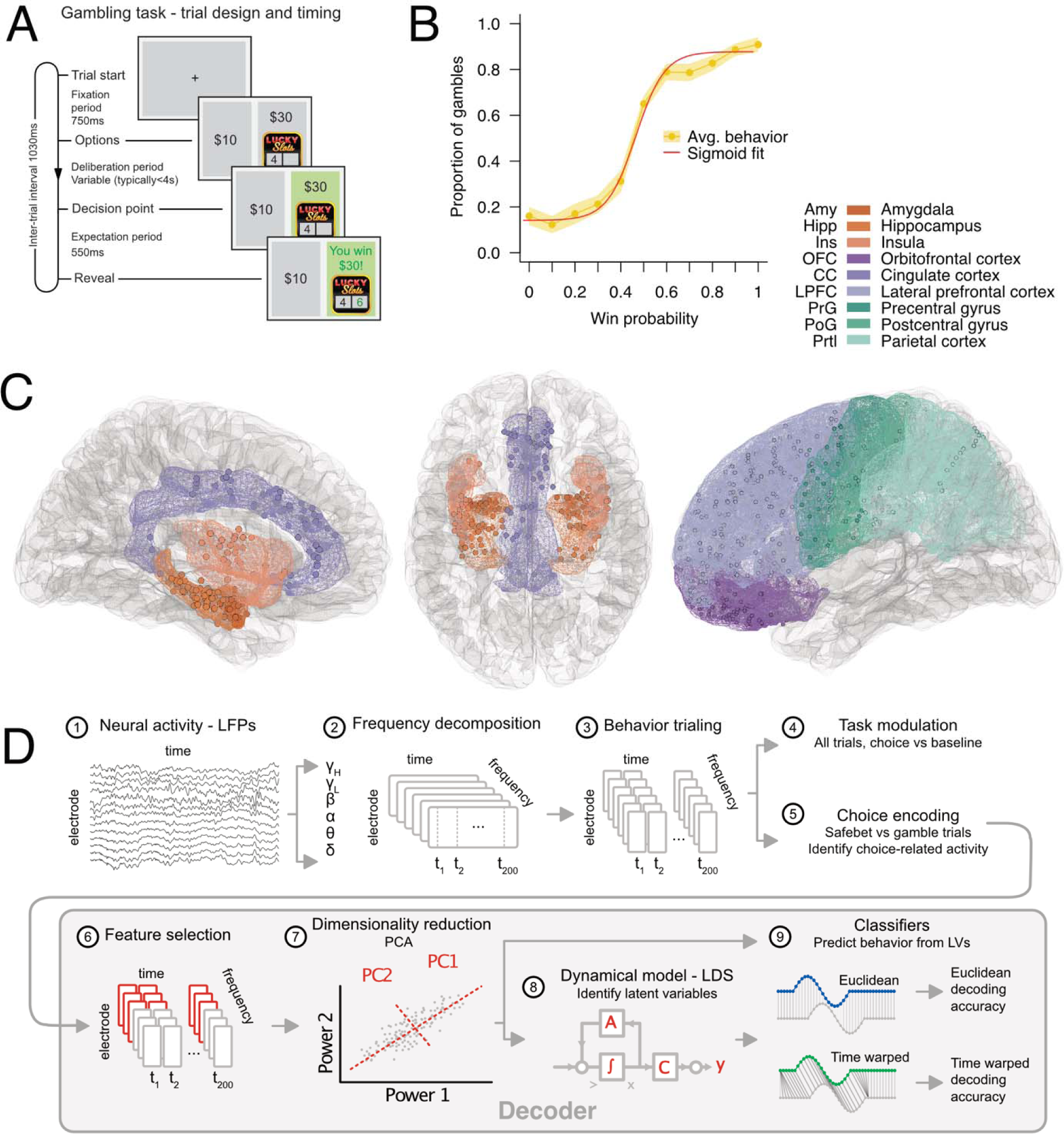
Experimental design and analytical strategy. a, Patients played a gambling game in which they chose between a safe $10 reward and a risky gamble for a higher reward. Gamble win probability varied parametrically on a trial-by-trial basis (0-100%). b, Average patient behavior (yellow line, shadow indicates SEM), shown as the proportion of risky choices (gambles) for each win probability, averaged in 10% increments. Patients gambled more often as the win probability increased. Red line represents a sigmoidal fit to the average behavior. c, Anatomical intracranial EEG (iEEG) coverage. We collected data from patients that underwent either electrocorticography (n=9, ECoG) or stereotactic EEG (n=11, sEEG) during epilepsy surgery (n=20 patients and 1042 electrodes total). We focused our analyses on electrodes located in grey matter in regions involved in reward-related behavior: orbitofrontal cortex (OFC), lateral prefrontal cortex (LPFC), precentral gyrus (PrG), cingulate cortex (CC), precentral gyrus (PrG), postcentral gyrus (PoG), parietal cortex (PC), amygdala (Amy), hippocampus (Hipp) and insula (Ins). d, Analytical strategy. Local field potentials (LFPs) from each grey matter electrode were cleaned and preprocessed (1), and subsequently decomposed into power across frequency bands (delta [1-4Hz], theta [4-8Hz], alpha [8-12Hz], beta [12-30Hz], gamma [30-70Hz] and high-frequency activity [HFA; 70-200Hz], 2). After trialing around behavioral events of interest (patient choice, 3), we determined which electrodes presented task modulation (significant power change versus baseline, 4) and choice activity (significantly different power between gamble and safe bet trials, 5) in any frequency band. Choice-active features (electrodes and frequencies) were selected (6) and subject to dimensionality reduction (7), followed by dynamical modeling and classification (8–9) or direct classification (9). We assessed model quality by examining the classification accuracy using a leave-one-out procedure.

### Power modulation patterns vary across regions

First, we assessed the electrophysiological nature of task-related neural activity during deliberation by examining power modulations in a 1-second epoch before choice. We analyzed local field potential (LFP) data from all electrodes located in grey matter in the above regions of interest (see Extended Data Table 1), and characterized power changes in each neural frequency band of interest (delta/δ [1-4Hz], theta/θ [4-8Hz], alpha/⍰ [8-12Hz], beta/β [12-30Hz], gamma/⍰] [30-70Hz], and high-frequency activity/HFA [70-200Hz]). We identified significant power modulations in each frequency band during choice by comparing the deliberation epoch (−1 to 0s before choice button press; see Methods, Extended Data Table 2 and Extended Data Figures 3 and 4 for reaction times and determination of optimal temporal window) to baseline (−0.5 to 0s prior to stimulus presentation) in each electrode. We found that 76.8% of electrodes (75.3%±0.25 across patients, mean±SD) showed significant modulation (either increase or decrease) at one or more frequency bands and were termed task-active (p < 0.01, paired t-test; Fig. 2a, Extended Data Tables 3 and 4). Electrodes were often task-active in multiple frequencies (2.09±1.70 frequency bands, mean±SD; Extended Data Table 5), with similar proportions of task-active electrodes across frequency bands (34.9%±0.05, mean±SD, Fig. 2b and Extended Data Table 6). There was variation in the proportion of task active electrodes, which ranged between 67.0% in OFC to 96.2% in parietal cortex (76.8±9.5, mean±SD, Fig. 2c, Extended Data Tables 3, 7 and 8), indicating widespread power modulation distributed across frequency bands and brain regions during deliberation.

**Figure 2.**
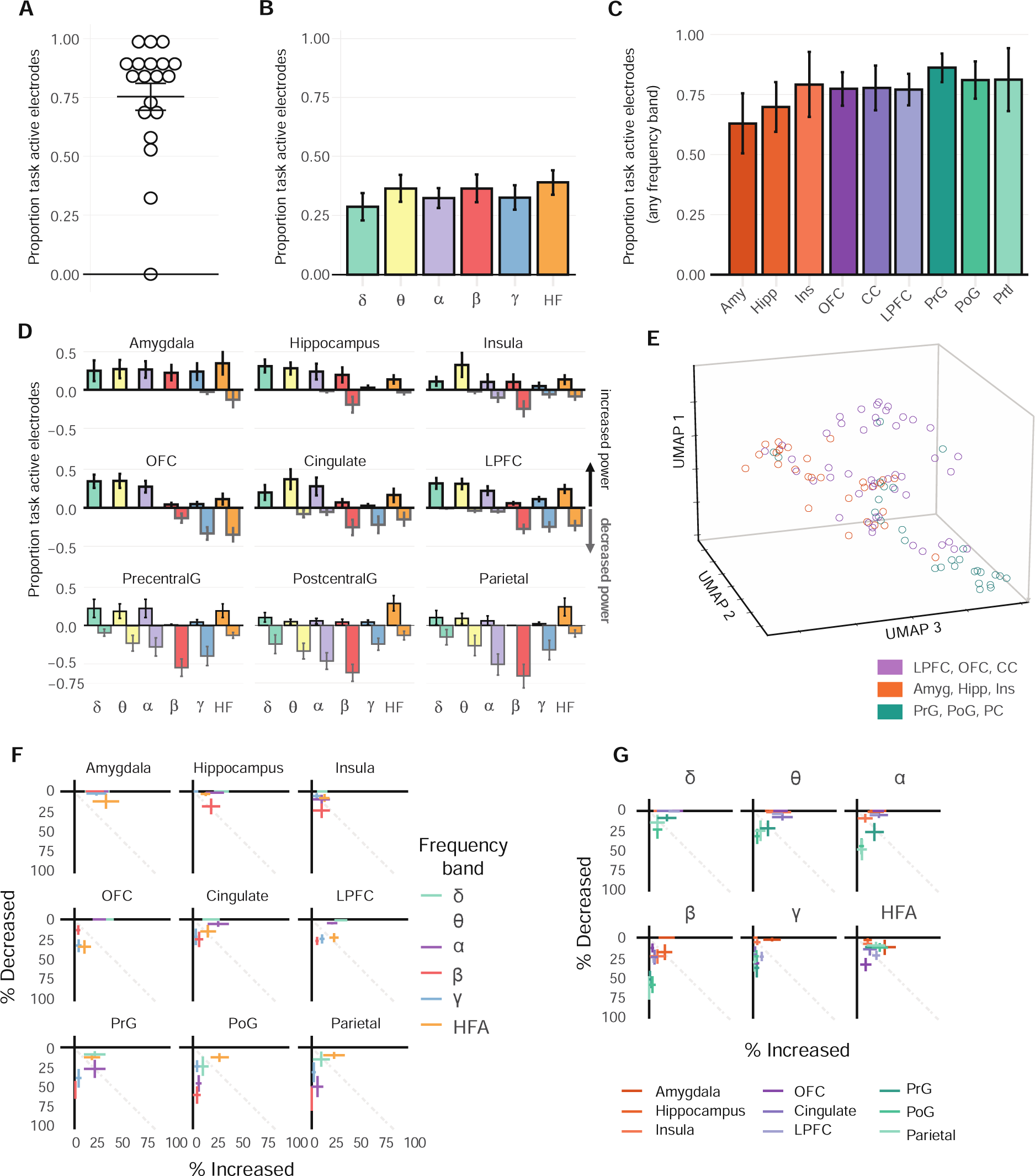
Proportion of task-active electrodes showing significant power modulation during deliberation. Power was compared between deliberation (−1s to 0s pre-choice) and baseline (fixation cross) epochs. a, Proportion of task active electrodes per patient. Horizontal lines show mean +/-SEM. b-c, Mean proportion of task-active electrodes across patients, grouped by frequency across all regions (b), and anatomical region across all frequencies (c). d, Mean proportion of task-active electrodes across patients, grouped by both region and frequency band. The proportion of electrodes that showed significant increases or decreases relative to baseline are shown as positive and negative bars on the vertical axis, respectively. e, Clustering of regions into functional groups: limbic (Amy, HC, Ins), prefrontal (OFC, LPFC, CC) and frontoparietal (PrG, PoG, PC). Clustered functional groups significantly above chance (bootstrapped p < 0.05) f, same data as in (c), but showing the patient-averaged proportion of electrodes showing increases (y-axis) or decreases (x-axis) separated by regions (panels) and frequency bands (points). g, Same data as in (c) but showing the patient-averaged proportion of electrodes showing increases (y-axis) or decreases (x-axis) separated by frequency bands (panels) and regions (points).

Because individual electrodes could show either increases or decreases in power, we separately quantified the proportion of electrodes showing power modulation in either direction. Overall, 50.5% of electrodes showed power increases and 55.6% showed decreases, with 29.3% of electrodes showing a combination of increases and decreases in separate frequency bands (e.g., an increase in delta accompanied by a decrease in beta; Extended Data Fig. 5). There was substantial heterogeneity in the proportion of electrodes that showed power increases/decreases across regions (Fig. 2d and Extended Data Fig. 5). For example, electrodes in Amy/HC predominantly showed power increases, whereas power decreases were most common in PC/PrG (Fig. 2d, 2f, and Extended Data Table 9). Grouping activations by frequency band instead of region shows a complementary depiction of power modulations. For example, power modulation in δ/θ consisted predominantly of power increases, whereas β/⍰] modulations consisted predominantly of power decreases (Fig. 2g, Extended Data Fig. 5 and Table 10). Finally, HFA was unique in that it showed bidirectional modulation in all regions, with individual electrodes within a region showing either increases or decreases (Fig. 2d, 2g, and Extended Data Fig. 5).

These observations show a rich pattern of region-frequency specific task-related power modulations, and revealed similarities among different groups of regions (Fig. 2d). For example, PoG/PrG/PC electrodes predominantly showed power decreases across most frequency bands (δ/θ/⍰/β/⍰]; average decrease = 35.5%); Amy/HC showed the opposite pattern, with widespread power increases across power bands (δ/θ/⍰/β/⍰]/HFA; average increase = 27%), whereas frontal regions (OFC/LPFC/CC), showed concomitant power increases in lower frequencies (δ/θ/⍰) and power decreases in higher frequencies (β/⍰]/HFA). Overall, we identified three putative sets of regions with similar modulation patterns: prefrontal (OFC/LPFC/CC), frontoparietal (PrG/PoG/PC) and limbic (Amy/HC/Ins) (Fig. 2e). To formally assess these similarities, we sought to classify regions according to their power modulation patterns using a clustering approach (see Methods). Briefly, we parameterized power modulation profiles by estimating the proportion of electrodes that showed increases or decreases in power modulation separately for each patient, region, and frequency band. Therefore, each patient-region combination (104 total data points, see Extended Data Table 1) is initially represented by a single point in 12D space (increase or decrease in power x 6 frequency bands) that represents its power modulation profile across frequency bands. After dimensionality reduction followed by classification using a k-nearest-neighbor algorithm, we identified three functional clusters with similar power modulation patterns, corresponding to prefrontal (OFC/LPFC/CC), frontoparietal (PrG/PoG/PC) and limbic (Amy/HC/Ins) circuits (Fig. 2e). To ensure the clustering algorithm was meaningfully separating each region into functional clusters, we carried out a permutation analysis (see Methods). The results further support the notion that patterns of power modulation are distinct across these proposed subcircuits (bootstrapped p < 0.05; Fig. 2e, Extended Data Table 11). In summary, the direction of power modulation (whether electrodes showed significant increases or decreases in power) depended on both frequency and region, defining three subcircuits whose power modulations were more similar within than between each subcircuit (prefrontal [OFC/LPFC/CC], frontoparietal [PrG/PoG/PC and limbic [Amy/HC/Ins]).

### Distributed high-frequency neural activity is related to choices

Because power modulations are likely to reflect multiple cognitive processes associated with choice behavior (e.g. sensorimotor, attentional)(27–29), the neural features associated specifically with choice were identified by assessing which electrodes showed a significant difference between safe bet and gamble trials in the 1 second prior to button press (choice-active electrodes; see Methods). Overall, 42.7% of electrodes were choice-active in one or multiple frequency bands (average = 0.55±0.73 frequency bands per electrode; Fig. 3a; Extended Data Tables 12 and 13). Choice-related activity was more common in higher frequencies (β/⍰]/HFA, mean = 6.05%/13.26%/19.72%, respectively) than in lower frequencies (δ/θ/⍰, mean = 2.76%/2.14%/3.24%, respectively; two-tailed t-test between low and high frequencies, p < 10^-6^; Fig. 3b, Extended Data Table 14). In contrast, the overall proportion of choice-active electrodes was largely homogeneous across anatomical regions (average = 37±7.7%), but greatest in frontoparietal regions (52.2%), followed by prefrontal regions (40.5%), and lastly limbic regions (33.1%) (see Fig. 3c, Extended Data Table 15). Moreover, we observed more choice-active electrodes in high-frequencies compared to low frequencies across all regions (Fig. 3d), suggesting a widespread and quantitatively comparable involvement of I] and HFA in choice behavior.

**Figure 3.**
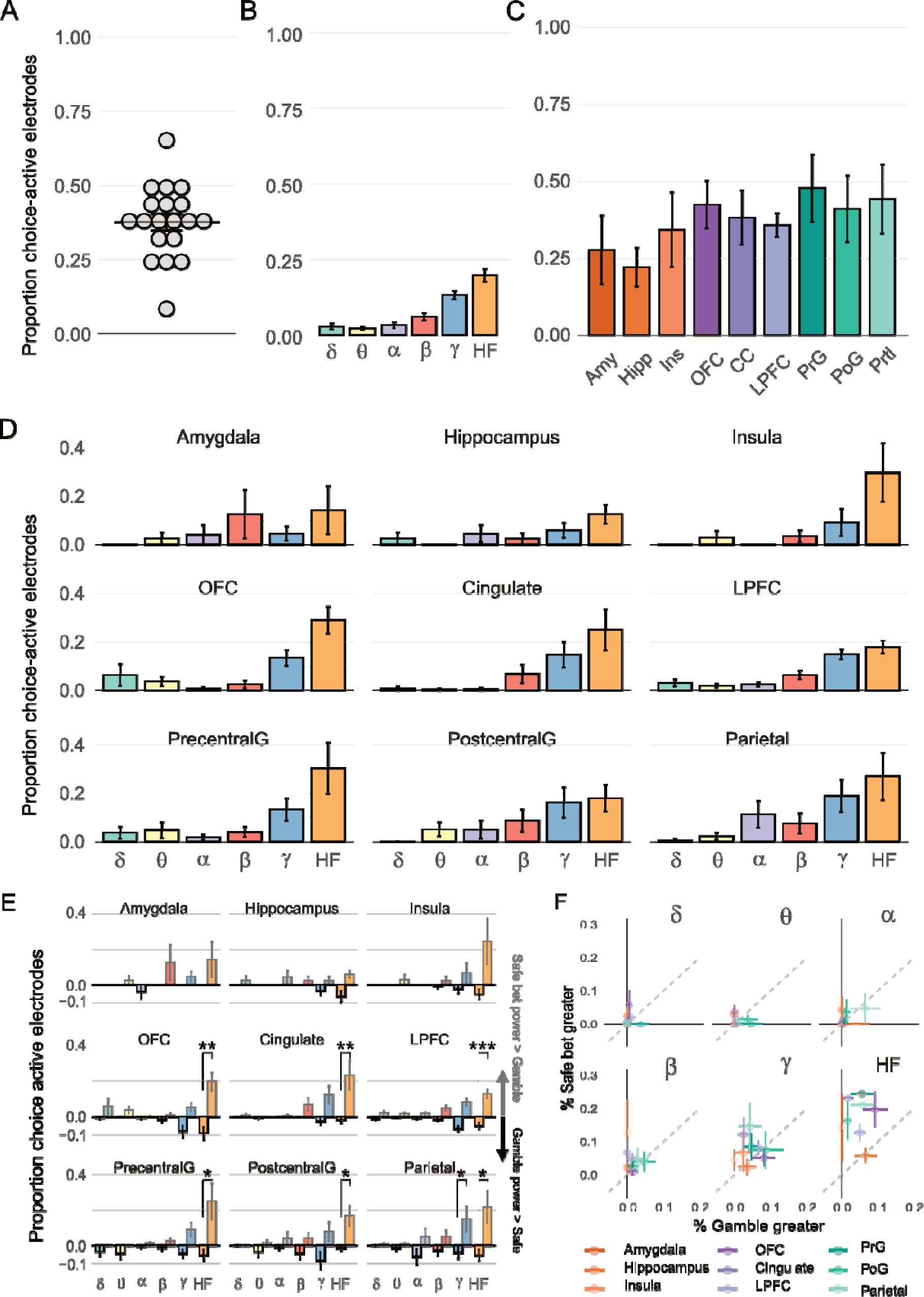
Proportion of choice-active electrodes showing a significant difference in power between safe bet and gamble trials. Power was compared between safe bet/gamble trials during deliberation (−1s to 0s pre-choice) using an analytical strategy that allowed differences in timing or intensity; An electrode was considered choice-active if there was a significant difference in power between gamble and safe bet trials in any frequency band (see Methods). a, Proportion of choice-active electrodes per patient (dots). Horizontal lines show mean +/-SEM. b-c, Mean proportion of choice-active electrodes across patients, grouped by frequency band (across all regions, b), and anatomical region (across all frequencies, c). d, Mean proportion of choice-active electrodes across subjects, grouped by both region and frequency. e, Proportion of electrodes separated by safe bet significantly greater than gamble (positive-going bars) and gamble greater than safe bet (negative bars). There were significantly more safe bet greater than gamble electrodes than vice versa (binomial test, p<0.001***, p<0.01**, p<0.05*). f, Same data as in e, grouped by power band (vertical axes, proportion safe bet greater; horizontal axes, proportion gamble greater).

Because HFA reflects local, non-rhythmic synaptic activity, we wanted to understand whether HFA power modulations were consistent across regions. Therefore, the relationship between gamble or safe bet choices and the relative increases or decreases in power across regions was examined. We observed significantly greater HFA power in safe bet compared to gamble choices than vice versa (Fig. 3e, f) in all regions examined except for hippocampus, amygdala and insula (binomial test, p < 0.05). This pattern was especially prominent for LPFC (p < 0.0001), Cingulate (p = 0.0012) and OFC (p = 0.0023). Taken together, these results indicate that differences in high-frequency power (I] and HFA) between choice/safe bet trials are widespread, appearing in most regions of interest, with safe bet trials associated with I]/HFA power increases compared to gamble trials.

### Power modulations encode multiple choice-related variables

Choice behavior depends on underlying reward-related computations, such as the win probability of the presented gamble, and other game features, such as the location of the chosen option on the screen. To probe the encoding of these choice-related variables across regions of interest the percentage of explained variance captured by the power modulations about the choice-related variables was used as an index of information encoding(17). Given the high proportion of HF choice-active electrodes shown above, and because HFA is known to reflect coordinated spiking activity among local populations of neurons(30–32), we focused on HFA for this analysis. For each choice-related variable, five behavioral regressors were identified: win probability, gamble risk, risky side, choice side, and choice (as above; see Methods, Extended Data Table 16 and Extended Data Fig. 6) using linear regression performed for each choice-related variable and time-point of the HFA time course, across trials. To determine whether neural activity in any given electrode encoded a specific regressor, we used a sum-of-F-statistic strategy that captures evidence in favor of HFA encoding across time points in combination with a permutation strategy (see Methods and Extended Data Table 17). After sorting the regressors by their ability to explain neural variance, we used a stepwise regression analysis to determine whether HFA activity reflected two or more regressors(17, 33) using a progressive model selection strategy in which a second (or third, etc.) behavioral regressor was added to the model only if there was a significant increase in explained neural variance (ANOVA p < 0.05).

We found that HFA activity encoded all of these choice-related variables including win probability (31±7% of all electrodes), risk (31±8%), risky side (33±9%), and choice side (33±10%, all p > 0.6, t-test), with a similar proportion of electrodes encoding choice-related variables across regions (Fig. 4). Choice was encoded in a significantly greater proportion of electrodes than all other regressors (48±8% mean±SD, Fig. 4a, paired t-tests vs other regressors all p < 0.01). Importantly, individual electrodes could encode multiple choice related variables. To quantify the degree of multiplexing, we compared the number of encoded regressors per electrode (Fig. 4b). We found that 21.2% (188/887) of all electrodes did not encode any regressors, 24.7% of all electrodes encoded a single regressor (219/887) and 54.1% (480/887) encoded two or more regressors (1.74±1.3 mean±SD regressors per electrode, Fig. 4b). The degree of multiplexing was similar across regions (Fig. 4c; ANOVA, p = 0.28), with 7/9 regions showing a mean between 1.50 and 2.00 encoded regressors per electrode (minimum in hippocampus, 1.34±1.11 and maximal in parietal cortex, 2.03±1.55). We observed differences in the extent to which individual regions represented each regressor (Fig. 4d, e). For example, risk was maximally encoded in OFC (47.2%), whereas choice side was maximally encoded in PoG (51.2%, see Extended Data Table 18). Choice, being most widely represented across areas, covers a larger area in the radar plot (Fig. 4d), compared to the other four regressors (t-test p < 0.01).

**Figure 4.**
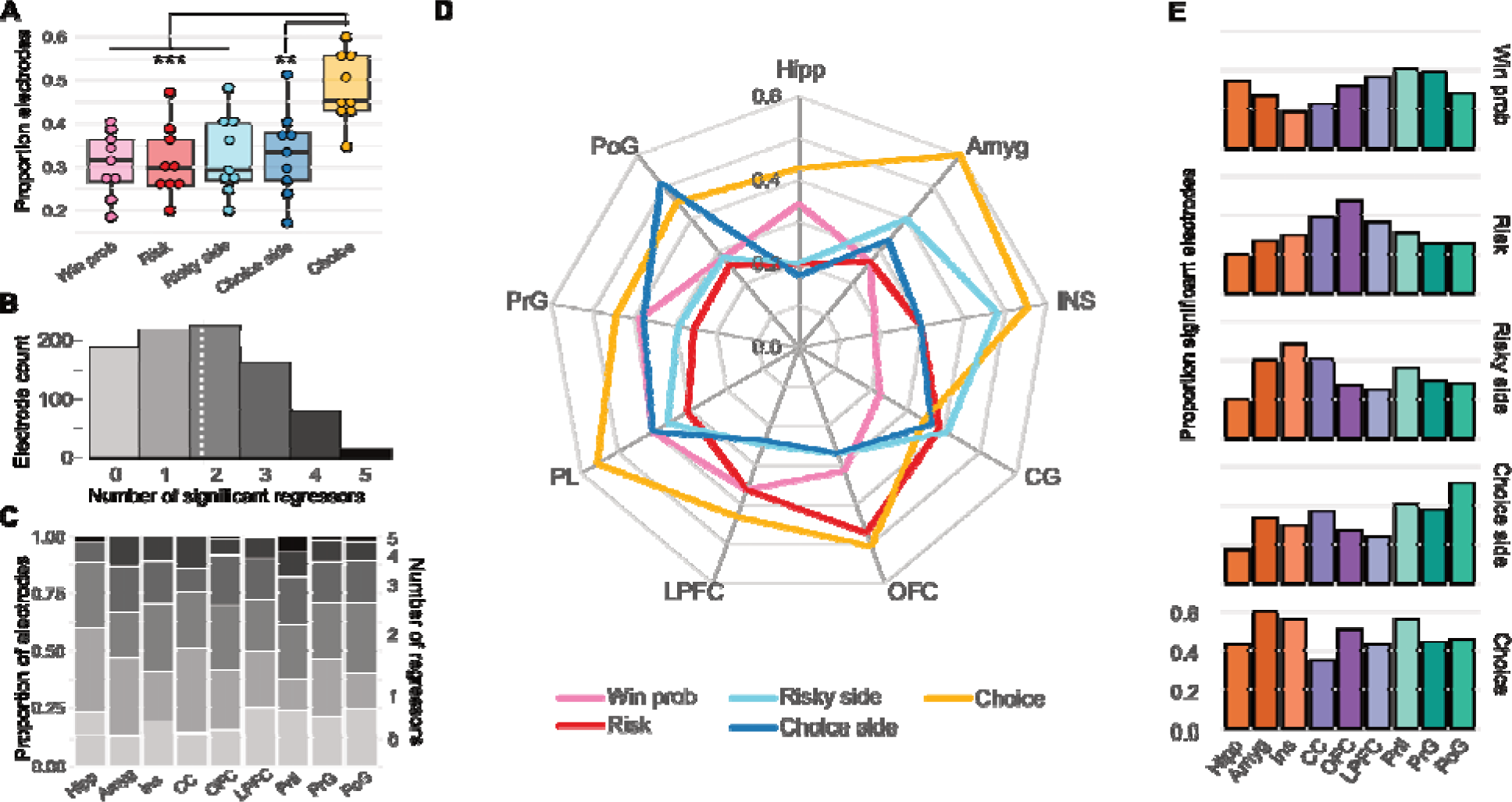
Counts and proportions of electrodes encoding choice-related regressors. a. Dot plots show average proportion of encoding electrodes per regressor (each dot corresponds to one region, averaged across patients); mean +/-s.e.m indicated by error bars. We examine encoding of win probability, risk, risky choice side (L/R), choice side (L/R), and choice (gamble/safe bet; see Text). Encoding for choice was significantly greater than all other regressors (paired t-test, p<0.005***, p<0.01**). b. Number of significant regressors encoded per electrode (regressors multiplexing): Histogram showing the number of electrodes that encode 0,1,2,3,4, or 5 regressors out of 5 regressors of interest. White dotted line indicates overall mean (1.74) c. Multiplexing: Proportion of electrodes per ROI with 0,1,2,3,4, or 5 significant regressors, gray stacked bars, shaded as in (a). There was no difference in mean number of significant regressors across regions (ANOVA, p=0.28). d. Radar plot shows proportion of significant electrodes per regressor (5 colored lines) per region (9 vertices). E. Bar plots show proportion of significant electrodes for each regressor and region of interest, grouped by regressor (same data as in D).

### Trial-by-trial choice can be decoded from high frequency activity

Next, we attempted to leverage the distributed nature of choice encoding to decode single-trial choices (e.g., risk vs safe bet) from neural activity alone. Since choice-active information is maximally present in I] and HFA (Fig. 3), we focused on activity in these frequency bands during the deliberation period (−1 s to 0 prior to button press) across all regions (see Methods and Extended Data Figure 4). To reduce the number of dimensions input to the decoder, we tested two dimensionality reduction strategies (principal component analysis [PCA] alone and linear dynamical systems [LDS] modeling after PCA) followed by two classifier strategies (a simple Euclidean distance classifier [ED](34) or dynamic time warping [DTW]). PCA and ED are standard methods for dimensionality reduction and classification, respectively; we chose to compare them to LDS and DTW which are potentially better suited to neural time series and may result in better decoding accuracy. LDS characterizes ongoing neural dynamics by identifying the underlying latent variables (LVs) that capture time-varying neural activity, and DTW is a distance metric that specifically accounts for temporal variation in neural activity (see Methods). Using a leave-one-out validation strategy and balanced accuracy, we found that the combination of LDS with DTW outperformed ED approaches (ANOVA F(4,19) = 39.1, p < 10^-17^; Fig. 5a). The optimal decoding strategy, LDS followed by DTW, accurately classified choice 74.3±3.4% across all patients and produced a maximal trial-by-trial single patient balanced decoding accuracy of 74.6%±3.19 (Extended Data Table 19). Therefore, a low dimensional, linear dynamical model followed by a classifier that accounted for variation in temporal dynamics across trials was the optimal strategy for choice decoding. Performance of the LDS+DTW was close to the maximal performance for all patients (Fig. 5a “Best”, 74.6%±3.2%, p > 0.01 Bonferroni corrected, paired t-test). Importantly, LDS+DTW achieved above-chance decoding in all patients, with a minimum decoding accuracy of 65.7%, indicating that LDS+DTW decoder was robust to variation in number of electrodes and anatomical localization across patients (Fig. 5a, Extended Data Table 19). In addition, we found that decoder performance depended on the win probability of the gamble trials, with the decoder performance falling to chance levels when win probability was either 0% or 100% (i.e., when there was no risk associated with the gamble choice). In contrast, decoder performance was similar for all risky choices (win probability between 10% and 90%; Fig. 5b, Extended Data Table 20). Therefore, we excluded trials with no uncertainty (gamble win probability = 0% or 100%) from decoder results (see Extended Data Tables 19 and 21 for comparison). Finally, we sought to examine the contribution of individual brain areas and subcircuits identified by task-related power modulation, which show different activation patterns (Fig. 2), to the decoding model. To do this, we carried out model training and classification as we progressively added (1) individual regions or (2) subcircuits (Fig. 2) to the decoding model (see Methods). For individual regions, we observed a linear increase in classification performance as we iteratively added the region with the least change in performance to the model. For subcircuits, we similarly found that increases in performance were not dependent on the specific subcircuit included (Extended Data Fig. 7). These results are also consistent with our results above that choice information is distributed across both individual brain regions and subcircuits (Fig. 3).

**Figure 5.**
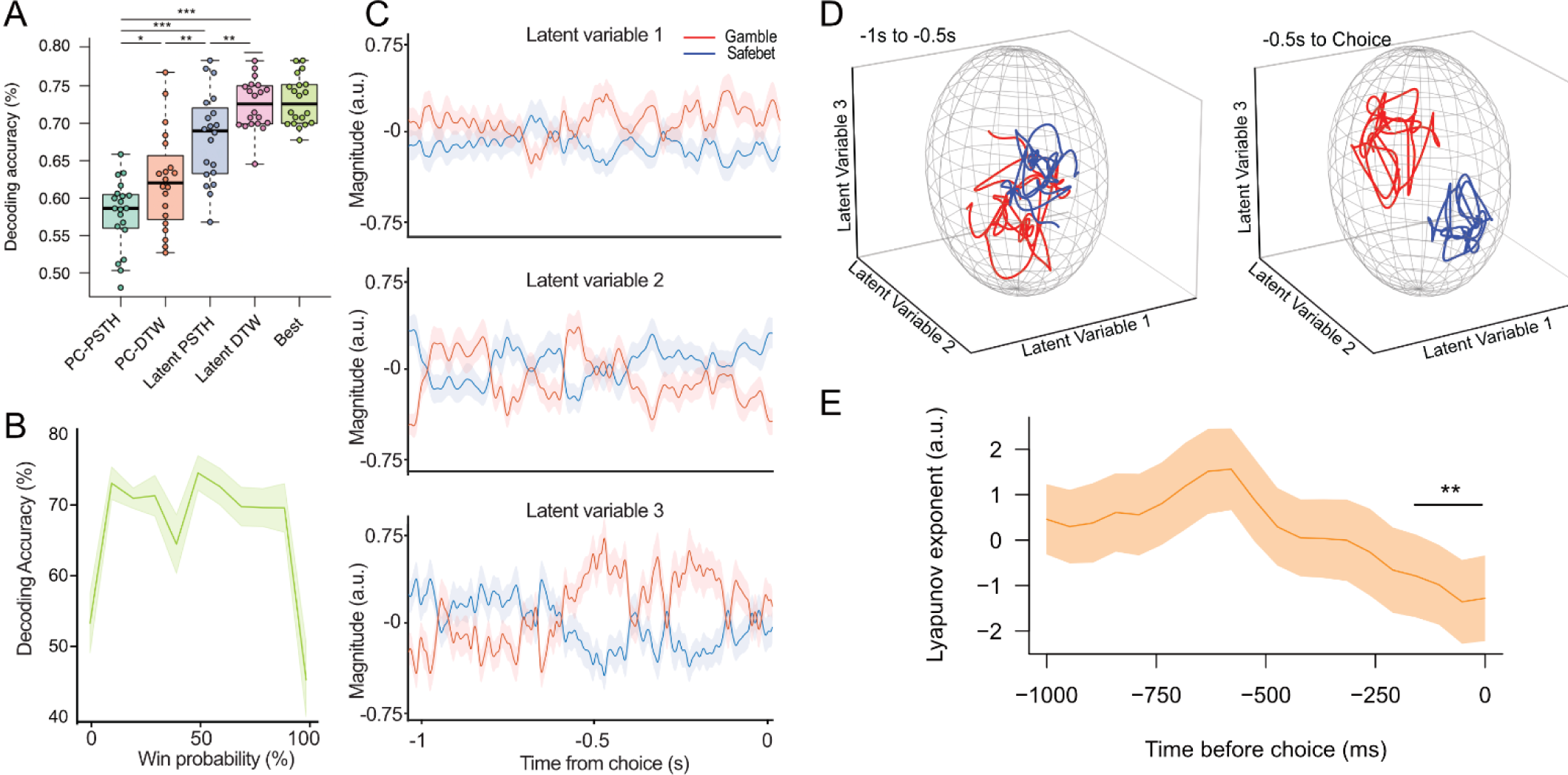
Choice decoding models and results. a, Classification performance for each decoding strategy, across all patients. We used two different dimensionality reduction methods, PCA and LDS, and two classification strategies, PSTH and DTW (see Methods). Results are shown for each combination of dimensionality reduction and classification. Latent DTW was significantly better than each other strategy (paired t-test, p<0.001***, p<0.01**, p<0.05*). The “best” category represents the highest balanced classification accuracy achieved for a given patient, across all strategies. b, Decoding accuracy for the LDS-DTW model as a function of trial win probability, averaged across all patients. c, Temporal evolution of LDS latent variables in an example patient. The plot shows the mean ± SE of the average latent variable trajectories for gamble (red) and safe bet (blue) trials. d, Latent variable trajectories in 3-dimensional space, separated in early (−1s to –0.5s pre-choice, top) and late (−0.5 to 0s pre-choice, bottom) deliberation period. e, Average Lyapunov exponents over time across subjects. Value goes from positive to negative at ∼-500ms indicating the trajectories becoming less chaotic and moving towards attractors with significant difference between –1000 to –600ms and –400 to 0ms (p<0.001, paired t-test).

### Latent variable modeling reveals sub-second neural dynamics underlying choice

Our LDS model defines a set of latent variables (LV) that describe the progression of neural dynamics through time. To gain further insight into the temporal evolution of neural activity during deliberation, we examined the evolution of LVs (Fig. 5c). Because some LVs may not reflect choice-related activity, we selected the three LVs whose representations during gamble trials were most different from safe bet trials (ED) and examined the LV projection for safe bet vs risky gamble choice trials (Fig. 5c and Methods). Each of these LVs therefore partially reflects the neural dynamic underlying choices. We observed that safe bet/gamble LV projections separate repeatedly during the deliberation phase, albeit with different temporal dynamics, especially within 500ms of choice (Fig. 5c). Thus, to gain an understanding of the temporal dynamics throughout deliberation, we examined the state space represented by the 3-dimensional trajectories defined by these 3 LVs as they unfolded in time (Fig. 5d). Early in the deliberation period (−1 s to −500 ms pre-choice), safe bet and gamble trajectories overlapped as the single trajectories moved back and forth from one end of the manifold to the other (Fig. 5d and Extended Data Movie 1). As the decision point approached (500 to 0ms pre-choice), safe bet and gamble LV trajectories diverged into non-overlapping subspaces of the state-space, suggesting the existence of separate attractors for gamble and safe bets (Fig. 5d). The generalizability of this individual observation is however limited by the fact that LVs are not comparable across patients. Other metrics such as classification accuracy (Fig. 5a) can be compared between patients, but they do not capture the evolution of neural activity changes over time. To address this issue, we estimated and compared Lyapunov exponents (see Methods), which represent whether a system is chaotic or converging to some attractors over time. Specifically, Lyapunov exponents measure the rate at which two initially similar multi-dimensional trajectories diverge from each other as time progresses(35) with positive values indicating that the trajectories are getting farther apart over time and negative values indicating the trajectories are getting closer together over time. Therefore, we calculated the Lyapunov exponent to measure the temporal rate of divergence separately for each subject and averaged them to examine temporal dynamics across patients. On average, we observed that the value of the exponents significantly shifted from positive to negative as the deliberation period approached the decision timepoint (p < 0.001 between −0.2 to 0s pre-decision compared to −1 s to 0.5 s prior to decision, paired t-test), indicating that the trajectories converged towards the attractors for gamble and safe bet as subjects neared a decision (Fig. 5e).

## Discussion

Value-based decision-making is well-known to depend on the coordinated activation of multiple brain areas, but multiple aspects of how neural activity across the human brain supports choices is not well understood, including the contribution of neural activity in different frequency bands, the distributed versus modular nature of choices and choice-related information, and the sub-second evolution of neural activity leading up to choices. Here, we sought to shed light onto these issues by leveraging surgical iEEG interventions to record distributed intracranial activity from prefrontal, fronto-parietal and subcortical reward-related regions during decision-making to examine the neurophysiological basis of human economic decisions. Our results show that although individual regions showed distinct neural frequency patterns of task-related power modulation (Fig. 2), choices were represented in high frequency activity (primarily I]/HFA; Fig. 3) across regions, in a highly distributed manner. We also observed encoding of choice-related information (win probability, risk, risky side, choice side), which was less distributed and showed higher regional modularity (Fig. 4). Moreover, latent variables capturing distributed neural activity reflected the sub-second dynamics underlying choices (Fig. 5). Despite differences in anatomical coverage and surgical strategy from patient to patient, these neural trajectories can robustly decode single trial activity to predict choice in every participant in our dataset (Fig. 5).

During the deliberation period, we observed power modulation across multiple frequency bands (Fig. 2), with rich patterns of bidirectional power modulation (increases/decreases) across regions. Whereas certain components of these task-related modulations are related to choice processes (see next section), multiple cognitive subprocesses engaged during decision-making tasks are reflected in oscillatory processes, including sensory processing(27), attention and target detection(29), and motor output(28). Consistent with this idea that oscillatory patterns across regions are functionally associated with different cognitive processes, we observed similar patterns of power modulation across functionally related brain regions, which our clustering analyses suggests correspond to subcircuits which we classified as limbic (amygdala, insula, hippocampus), prefrontal (LPFC, OFC, CC) and fronto-parietal (PrG, PoG, Prtl) networks (Fig. 2d). One possibility is that these common low-frequency power modulations reflect the neural basis of individual cognitive processes engaged in decision-making, whose activity profiles are likely to vary across the areas in these subcircuits according to their individual involvement. For example, we observed widespread beta-band power decreases in frontoparietal contacts (Fig. 2d), possibly reflecting beta desynchronization associated with pre-motor processes(36). δ-θ increases were widespread in limbic and prefrontal electrodes, an observation consistent with the appearance of θ-band oscillations in the hippocampus and prefrontal cortex during goal-directed behavior, navigation and memory formation(37, 38). Attentional processes in turn engage oscillations in the I] and I] frequency bands(39, 40). Therefore, one possibility is that the complex power modulation patterns we observe reflect the simultaneous activation of these different processes, an idea supported by our observations of common patterns of power modulation across anatomically and functionally related brain regions (Fig. 4). Alternatively, similar patterns of low-frequency power modulation may reflect the establishment of functional communication across regions, in a manner reminiscent of “spectral fingerprinting” in which frequency-specific activations reflect computations associated with specific cognitive processes and interactions among regions(41). Finally, task-related HFA modulation was bidirectional (Fig. 2), reflecting heterogeneity in local spike rate encoding(24, 42, 43) that may be modulated by changes in correlated activity of local populations of neurons(30, 44).

Compared to this heterogeneity in task-related activation across regions, we observed a striking similarity in choice-encoding across regions, with activity in higher frequencies (I]/HFA) discriminating whether the patient opted for the safe/risky choice in all regions in our sample (Figs. 3 and 4). The importance of I]/HFA in choice encoding is consistent with the notion that broadband I] activity reflects local neuronal activation related to value-based decision-making(17, 18, 24, 42, 45, 46). Choice activity was present in I]/HFA (Fig. 3), encoded to a high degree in HFA (Fig. 4), and was decodable from all regions studied (Fig. 5), indicating widespread representation of choice information. Interestingly, unlike power modulations, which were often bidirectional (increases/decreases), we observed a consistent relationship between high-frequency activation and the quality of the choice, such that high-frequency activity was higher in safe bet than risky choice trials (Fig, 3e-f), suggesting that selecting the safe option was consistently associated with a higher degree of neuronal activation. This pattern was especially prominent in LPFC, cingulate cortex, and OFC, which suggests exertion of cognitive control to obtain a potentially lower, but safer, reward(47–49).

The ubiquity of choice-related I]/HFA encoding across areas in our study is consistent with the view that value-based decision-making is a distributed process engaging multiple brain areas, likely simultaneously. The growing consensus on the existence of widespread brain activation for multiple cognitive processes including decision-making is supported by an increasing amount of evidence from primate single unit recordings(4, 5, 49–51), rodent(6, 7), and human imaging studies(52). Our data support a distributed model of decision-making(50, 53), which extends ‘action-plan’ models(54–56) by proposing that decision-making is a parallel, distributed process that weighs information across all representational levels, including those of good-based valuation and up until action. Importantly, the observation that choice is widely represented by high frequency activity across all brain areas studied is not inconsistent with our observation of different power modulation profiles across regions (see above); rather it supports the notion that decision processes only partially drive power modulations, which are likely to also reflect a variety of other sensory, cognitive, and premotor processes.

Decisions involve several underlying computations related to both the utility of the presented choices as well as the sensorimotor aspects of the task (i.e., choice locations and motor plans associated with choice) which are known to be present in different brain areas. Here, we examined a variety of computations associated with trial-by-trial choices, including the win probability(57), risk(58), spatial location of the presented options and motor command to express the selected choice. We observed widespread encoding of these variables (Fig. 4), similar to choice information albeit with two important differences: first, choice information engaged a significantly higher proportion of electrodes across brain areas (Fig. 4c) than any of these computations, and second, there were specific modular profiles – not all computations were equally represented in all regions (Fig. 4d). For example, risk information was significantly more represented in OFC than in any other region studied, consistent with previous fMRI(58, 59) and electrophysiological(17) evidence; motor-associated activity (left/right choice), which was orthogonal by design to the quality of the choice (safe/risky) was primarily represented in motor and premotor areas (PrG, PoG); encoding of the risky side was predominant in insula, amygdala and CG regions, consistent with their role in arousal, anticipation and conflict or error monitoring(49, 60, 61). These observations are consistent with more modular, regionalized decision computations that coalesce into a more salient choice signal that is more widely represented across regions. Conceptually, win probability and risk are more related to a good-based space(2, 33), since they represent modifiers of the quality of the gamble choice (and therefore modulate value); risky side and choice side relate to the visuo-motor aspects of the task, and therefore more closely related to action-based space. The observations that these computations are unevenly distributed across brain areas are consistent with good-based models which predict functional separation of valuation and spatiomotor information(2); however, despite this unequal representation, all regressors were encoded in a substantial proportion of electrodes across regions (35.1±10.5%, mean±SD across all regions), consistent with distributed consensus models that predict that choice representations compete simultaneously and flexibly right until the moment that the action is executed(50, 53). Therefore, our observations propose an intermediate implementation whereby computations subserving choice are distributed across multiple decision-related areas, but with a certain degree of regional specification.

Using a low dimensional state space model, we were able to accurately classify trial-by-trial choices with high accuracy (74.3±3.4%). This decoding model performed significantly above chance in all patients in our sample (Fig. 5a), despite substantial differences in electrode location (Extended Data Fig. 2). This robust decoding performance is likely due to the highly distributed nature of choice information, which meant that a substantial proportion of electrodes reflected choice across all regions studied (Fig. 3) and therefore the model could perform regardless of variation in electrode number and anatomical location. This result suggests that decoding models reach higher accuracy performances using distributed recordings, rather than a more limited set of ROIs. More generally, these results demonstrate that distributed decoding models are capable of identifying abstract information, adding to previous studies showing successful decoding of mental states from iEEG data(62–64) and demonstrate the possibility of developing robust decoders for behavioral data. Importantly, time warping algorithms (DTW) resulted in the best average decoding performance across tested models, suggesting that decision-encoding across sites and patients had consistent yet time-varying temporal dynamics that impact the ability of models to decode choices.

Our dynamical systems model generated low-dimensional representations (latent variables) of neural activity, which traversed the state space during deliberation before settling on one of two attractors, for either safe bet or gamble decisions (Fig. 5). Interestingly, individual LVs showed rapid switching (at approximately 1-4 Hz) during deliberation, suggesting that both safe bet/gamble attractors were visited multiple times before a choice was made (Fig. 5c). This is similar to activity patterns observed during deliberation in multi-electrode OFC decoding in non-human primates that reflect the fast alternative evaluation of binary choices as in our task(45, 65). Moreover, the 3-dimensional trajectories of the 3 LVs that best decoded choice revealed a progressive separation of gamble and safe bet trajectories (Fig. 5d), suggesting that deliberation in this higher-dimensional space may evolve progressively until a decision-threshold is reached(66, 67). Therefore, our results suggest that, during the deliberative process both rapid switching and evidence accumulation occur simultaneously. Rapid switching can be observed when examining the trajectory of a single LV across time, whereas representations in higher dimensional space (multiple LVs) reveal progressive separation of the neural dynamics, suggesting accumulation of information towards the final choice attractor.

Finally, our decoder could not decode non-ambiguous ‘catch’ trials (win probability equal to 0% or 100%) where risk was zero and therefore not a factor for choice. This suggests that the model was appropriately sensitive to conditions where deliberations based on risk evaluation were used to make the choice. In contrast, performance across a broad range of win probabilities was similar, suggesting that the neural representation of these similar computations is correspondingly similar.

Our results provide several novel observations on the neurobiological basis of human decision-making. First, we show a differential impact of activity in higher (β/⍰]/HFA), but not lower frequency bands, containing information about the upcoming choice during deliberation. As HFA is highly correlated with fMRI BOLD signal(30, 44), this observation directly implicates neuronal activity in choice, and helps bridge fMRI studies with animal studies that implicate SUA or MUA activity, which is also reflected in high-frequency LFP activity(24, 42, 44), in decision-making processes. Second, we demonstrate a highly distributed nature of human decision-making processes across regions, consistent with observations in animal models(6, 7, 50, 53), while providing evidence for a more modular encoding of several decision-related variables such as risk(58), win probability(57) and motor activity(28) consistent with observations from non-invasive studies. Given their highly distributed nature, our sEEG recordings cover a wide number of areas and provide a more complete depiction of electrophysiological activity than in most animal model studies which are limited to a few areas(68, 69). Third, we show the sub-second temporal dynamics of decision processes, which develop in the ∼500ms prior to choice but not earlier. Finally, we achieve a significantly higher decoding accuracy (74.6%±3.19%) than comparable non-invasive decoding approaches. For example, Vickery et al (2011) used MVPA whole-brain approaches to decode both outcome (win/loss) information and choice (stay/shift) from distributed brain activity, but only achieved a mean accuracy of 52.6% for binary choices. A similar study using voxel-based techniques, found decoding accuracies as high as 64% in ACC(57). A recent (2018) review reported that typical fMRI MVPA-based decoding accuracies for reward-related signals are a few percent points above chance(70), suggesting that intracranial EEG methods provide a much more sensitive measure for decoding choices.

In summary, here we combined invasive iEEG recordings, economic probes of decision-making, and machine learning approaches to characterize circuit-wide activity underlying decision-making behavior with high anatomical specificity, temporal resolution, and neurobiological detail. Our results show that (1) deliberations are associated with power modulation patterns that are different across functional clusters of regions (limbic, prefrontal, fronto-parietal); (2) choice information is anatomically distributed, present in high (I]/HFA) frequency activity in a set of cortical and subcortical regions; (3) individual computations related to choice are also widespread, but overall less strongly represented and show a certain amount of regional specificity; and that (4) high-accuracy (74.6%) decoding of trial-by-trial behavior from sEEG data is possible and robust to variation in electrode placement across patients. Therefore, our study provides support for the view that highly distributed patterns of high-frequency activity support economic choices, which in turn allows for high-accuracy, robust trial-by-trial decoding of risky decisions from intracranial data. Future efforts will be tasked with development of general but patient-individualized brain-state decoders for other cognitive (i.e., memory, attention) and translational (i.e., pathological vs healthy brain states) states from distributed sEEG activity alone. Combined with the ability to directly and precisely modulate brain activity using direct electrical stimulation(71–78), these results open the door to the development of cognitive prostheses to modulate abstract brain states in the human brain.

## Acknowledgements

We would like to thank L. Nuñez, C. Meikle, and C. Foreman for help with data collection. We would especially like to thank the patients for their willingness to participate in this research. The project described was supported by the National Institute of Mental Health through grant number K01MH108815, and the National Center for Advancing Translational Sciences, National Institutes of Health, through grant number UL1 TR001860 and linked award TL1 TR001861. The content is solely the responsibility of the authors and does not necessarily represent the official views of the NIH. The authors declare no competing interests.

## Methods

### Subjects

Data were collected from 34 patients (19 female) with intractable epilepsy who were implanted with chronic subdural grid or strip electrodes (electrocorticography, ECoG) or stereotactic EEG (sEEG) electrodes as part of a procedure to localize the epileptogenic focus. Electrode placement was based solely on the clinical needs of each patient. Data were recorded postoperatively in the epilepsy monitoring unit at five hospitals: The University of California (UC), San Francisco Hospital (n = 3), the Stanford School of Medicine (n = 3), UC Irvine Medical Center (n = 23), and UC Davis Medical Center (n = 5). As part of the clinical observation procedure, patients were off anti-epileptic medication during these experiments. All participants gave written informed consent to participate in the study in accordance with the University of California, Davis or University of California, Berkeley Institutional Review Board. Patients understood that they could decline participation at any time, and verbal assent was reaffirmed prior to each experimental task.

### Electrophysiological data acquisition

ECoG and sEEG activity was recorded, deidentified, and stored at the same time as behavioral data. Data were collected using Tucker-Davis Technologies, Nihon-Kohden, or Natus systems. Data processing was identical across all sites: channels were amplified x10000, analog filtered (0.01 - 1000 Hz) with > 1kHz digitization rate, re-referenced to a common average offline, high-pass filtered at 1.0 Hz with a symmetrical (phase true) finite impulse response (FIR) filter (∼35 dB/octave roll-off). Behavioral data were simultaneously collected using a PC laptop running Python (v.2.7) and PsychoPy (v.1.85.2) and synchronized with a timed visual stimulus (trial start) recorded by a photodiode through an analog input to the electrophysiological system.

### Behavioral task

We probed risk-reward tradeoffs using a simple gambling task described previously(17). Briefly, patients chose between a safe bet ($10, fixed) or a gamble for potential higher winnings (between $15 and $30). Gamble win probability varied per trial based on an integer between 0-10 shown at game presentation. At the time of outcome (t = 550ms post-choice, Fig. 1a), a second number (also 0 - 10) is revealed. The gamble results in a win if the second number is greater than the first, and ties were not allowed, therefore, a shown ‘2’ had a win probability of 80% and an ‘8’ had a win probability of 20%. Both numbers were randomly generated using a uniform distribution. Location of safe bet and gamble options (left/right) were randomized across trials. Patients played 10 practice trials, repeated as many times as necessary, to ensure they had full knowledge of the (fair) structure of the task prior to game play (200 trials). Timing is summarized in Figure 1a. Trials started with a fixation cross (t = 0), followed by a game presentation screen (t = 750ms). Patients had up to 8s to respond (mean reaction time = 1.4s). Gamble outcome presentation appeared 550ms after button press (choice) on each trial regardless of choice. A new round started 1s after outcome reveal. The experimental task typically lasted 12-15min. This gambling task minimized other cognitive demands (working memory, learning, etc.) on our participants while allowing us to test for decision-making under risk. Behavioral performance was assessed by examining the proportion of trials in which the patient chose to gamble as a function of win probability; the proportion of risky trials was calculated for each win probability value (0-100% in 10% increments) and fit with a logistic curve (Fig. 1B, Extended Data Fig. 1). As a control for behavioral data quality, we excluded patients in which a logistic function did not appropriately fit the relationship between percentage of gambles and win probability (p<0.05, logistic fit). Results from a subset of these patients playing the same task were published previously(17). Fourteen (14) patients did not show a significant fit and were removed from further analysis, leaving twenty (n = 20) subjects with behavioral data of sufficient quality. This high exclusion rate largely reflects the demands of data collection with patients in a clinical environment as well as the importance of stringent behavioral criteria.

### Anatomical analyses

Electrode localization was based strictly on clinical criteria for each patient, 9/20 had electrocorticography (ECoG) grids, predominantly in orbitofrontal, lateral prefrontal, and parietal regions, whereas 11/20 had stereotactic EEG (sEEG) coverage, predominantly of deep temporal lobe regions (amygdala, hippocampus) (Extended Data Fig. 2). For each patient, we collected a pre-operative anatomical MRI (T1) image and a post-implantation CT scan. The CT scan allows identification of individual electrodes but offers poor anatomical resolution, making it difficult to determine their anatomical location. Therefore, the CT scan was realigned to the pre-operative MRI scan following a previously described procedure(79). Briefly, both the MRI and CT images were aligned to a common coordinate system and fused with each other using a rigid body transformation. Following CT-MR co-registration, we compensated for brain shift, an inward sinking and shrinking of brain tissue caused by the implantation surgery. A hull of the patient brain was generated using the FreeSurfer analysis suite, and each grid and strip was realigned independently onto the hull. This step was necessary to avoid localization errors of several millimeters common in ECoG patients. Subsequently, each patient’s brain and the corresponding electrode locations were normalized to a template using a volume-based normalization technique and snapped to the cortical surface(79). Finally, the electrode coordinates are cross-referenced with labeled anatomical atlases (Brainnetome atlas) to obtain the gross anatomical location of the electrodes, verified by visual confirmation of electrode location based on surgical notes. We selected all grey matter electrodes across a broad set of regions known to be involved in reward-related behavior for analysis (Fig. 1C): lateral prefrontal cortex (LPFC; 391 electrodes from n=19 patients), orbitofrontal cortex (OFC; 193, n=18), cingulate cortex (CC, 84, n=13), hippocampus (HC; 65, n=13), amygdala (Amy; 32, n=11), insula (Ins; 46, n=8), precentral gyrus (PrG; 108, n=11), postcentral gyrus (PoG; 88, n=9), and parietal cortex (PC; 78, n=8) (Fig. 1; see Extended Data Table 1 for a complete account of electrode numbers across regions and patients).

### Electrophysiological analyses

#### Quality control and preprocessing

Epileptogenic channels and channels with excessive noise (low signal-to-noise ratio, 60 Hz line interference, electromagnetic equipment noise, amplifier saturation, poor contact with cortical surface) were identified and deleted. Out of 1194 electrodes localized to regions of interest, 1085 were artifact-free and included in subsequent analyses. Additionally, all channels were visually inspected to exclude epochs of aberrant or noisy activity (typically <1% of datapoints). Data analysis was carried using custom scripts written in MATLAB and Fieldtrip toolbox(80). Data for each channel was downsampled to 1KHz. each channel was lowpass filtered (200Hz), highpass filtered (1Hz), and notch filtered (60Hz and harmonics) to remove line noise, and downsampled to 1 kHz if necessary. Electrode channels were re-referenced to a common average reference of all electrodes in each strip/grid. Even though bipolar derivations or white matter referencing are often used for sEEG electrodes, we opted to use a single (CAR) re-referencing strategy for both ECoG and sEEG electrodes for analytical consistency. Trials were epoched to the time of decision using a [-4,3]s window around events of interest (options presentation, and patient choice), and the leading and trailing 1s of data were discarded to remove edge effects. Time-frequency representations (DPSS taper method) were plotted for each region and patient (averaged across electrodes and trials) and visually inspected for artifacts.

#### Time-frequency representation of neural activity (bandpass estimates)

To examine the role of individual oscillatory bands, we decomposed the neural activity into canonical, discrete activity bands: (delta, δ [1-4Hz]; theta, θ [4-8Hz]; alpha, α [8-12Hz]; beta, β [12-30Hz]; gamma, I] [30-70Hz]; high frequency activity, HFA [70-200Hz]) for each grey matter electrode for each patient using the Filter-Hilbert method. Power in the 6 bands was calculated by applying a Butterworth bandpass filter (order 3 for delta and order 4 for all other power bands) and Hilbert transform and multiplying the resultant complex signal by its complex conjugate(81). As before, one second is removed from the beginning and end of the data to reduce edge effects. Prior to dimensionality reduction for classification, power data for each trial and channel was smoothed and downsampled using a 50ms sliding window with 10ms step increments(17) and z-scored over the time dimension within each band to correct for the 1/f profile of neural activity.

#### Task-Active Electrodes

To examine patterns of electrodes that showed significant task-related power modulation, we compared average power estimates during deliberation period (−1 to 0s pre-choice) with baseline power estimates (−0.5 to 0 pre-stimulus onset) for each electrode and each frequency band independently (paired t-test across trials, alpha = 0.01). Deliberation period of −1 to 0s pre-choice (button press) was determined using a forward and backward classifier temporal window analysis (Extended Data Figure 4). Data were analyzed and plotted using custom scripts in Matlab and R. To investigate patterns across patients, regions, and powerbands, we summarized and plotted the proportion of task-active electrodes for each patient (Fig. 2A, Extended Data Tables 3), and then calculated the mean proportion of task-active electrodes across patients for each power band (Fig. 2b, Extended Data Table 5) and region (Fig. 2c, Extended Data Table 7). Mean patient proportions and standard errors provide a depiction of population activity across patients that would be obscured in the aggregate. For comparison, proportions of task modulated electrodes overall (n=1042 total electrodes) are summarized separately (Extended Data Tables 2,4,6). Finally, in order to probe the apparent homogeneity of the results, we plotted proportions of task active electrodes that increased or decreased power per power band and regions (Fig. 2d, f, g, Extended Data Figure 3, and Tables 8,9) which revealed patterns that were quantified with clustering methods below.

#### Region Clustering

We applied an unsupervised clustering algorithm (k-nearest neighbor) to evaluate the degree of similarity in power modulations across regions. We started by parameterizing activity in each brain region by estimating the proportion of electrodes that showed increases or decreases in power modulation during the deliberation period, separately for each patient, region, and frequency band (see Task Modulation, above). Therefore, each patient-region combination (104 total data points, see Extended Data Table 1) is initially represented by a single point in 12D space (increase or decrease in power x 6 frequency bands). To reduce noise or redundancy, we performed dimensionality reduction using Uniform Manifold Approximation and Projection(82). UMAP converts the data into a k-neighbor graph and then identifies a projection into a lower dimensional space by minimizing the cross-entropy between the two representations while maintaining the fundamental characteristics of the original graph. Once the data were projected into a lower dimension we applied the k-nearest-neighbor algorithm to sort the data into a specified number of clusters. In order to ensure the clustering algorithm was meaningfully separating regions into super region clusters, we bootstrapped our data by repeatedly (1000 times) randomly shuffling the labels on each region and rerunning the clustering algorithm to tested the likelihood that regions would be randomly separated into super region clusters. We used that distribution to evaluate the significance of each cluster. This method effectively separated the regions into three hypothesized functional groups (bootstrapped p < .05) (Extended Data Table 9).

#### Choice-Active Electrodes

Next, we defined choice-active electrodes as those that showed a significant period of activation during deliberation (−1 to 0s pre-choice) between trials where a gamble bet was chosen compared to safe bet. This was determined by permutation test. Trials were separated by subsequent choice (gamble or safe bet) and then a two-sample t-test was applied to every time point (1 ms resolution) in the 1-second deliberation epoch. MATLAB’s `bwconnect` function was used to find contiguous suprathreshold clusters of points (alpha < 0.05). An electrode was categorized as choice-active by comparing the sum of the T-statistic for the largest cluster to a trial-shuffled null distribution at alpha of 0.001 (2-tailed). To determine the null distribution, trial labels were shuffled 10,000 times and the sum of the T-statistic of the largest supratheshold cluster was recorded on each iteration. This was repeated for each electrode, for each power band, for each patient. Similar to task-active electrodes, we plotted the mean proportion of choice-active electrodes for each patient (Fig. 3A, Extended Data Table 11), and summarized mean proportions of choice-active electrodes across patients by frequency band (Fig. 3B, Extended Data Table 14) and region (Fig. 3c, Extended Data Table 13), as well as separated by region and frequency (Fig. 3d), and separating by whether power was greater for safe bet or risky gamble choice (Fig3e, f). Regions and frequency bands with the greatest proportions of choice active were selected as features for subsequent choice decoding (below). Patients with less than 50 trials after artifact rejection were excluded from these analyses, as well as regression analyses below.

#### Electrode-wise regression analysis

In order to identify which behavioral and computational regressors were reflected in high frequency activity (HFA) [60-200Hz] in each of nine regions of interest we performed a stepwise regression analysis on an electrode-by-electrode basis. Data from 14 patients were included, as 6/20 patients had low trial number counts and were excluded from this analysis and decoding (see Supplementary Methods Table 15 for summary of electrode counts patient and region). As described for other analysis, power-time vectors for each trial were smoother using a rolling window average (200ms window width, 50ms step increments). The resulting trials x power (over time) matrix was used to carry out stepwise regression as described below.

#### Regressors of interest

We chose a set of 5 regressors related to decision/choice and action, which showed low to moderate collinearity (Supplementary Figure 5). The 5 remaining regressors were win probability, risk, risky side index, gamble index, and buttonpress index. Briefly, win probability represented the probability the gamble option resulting in a win (winprob = (10 – number shown)/10); risk represents the level of uncertainty associated with a gamble (Risk = (winprob .* (1-winprob)); 0 if winprob is 0 or 1, and maximal if winprob = 0.5); risky side is a Boolean regressor indicating which side the risky side is presented on (right =1); choice side is a Boolean regressor indicating which side was chosen (right = 1) and choice represents the patient choice as above (1=gamble, 0=safe bet).

#### Stepwise Regression Strategy

As an initial step, and to reduce processing time, we ran independent linear regressions for each electrode for each regressor of interest and each time point; regressors that were significant at this stage (p < 0.05) were passed to a sum of F-statistic permutation test (1000 iterations; see next section for details) and the resulting p-value was multiple comparisons corrected over the number of electrodes in our sample (n=978). The regressors that survived permutation and multiple comparisons were then passed to a stepwise regression model. The best regressor order was selected by starting with the regressor with the highest association (R^2^), and then adding each possible subsequent regressor. The model that explained the most variance was chosen based on model comparison of the reduced model (reg_1_) to the complex model (reg_1_ + reg_2_) using an ANOVA test. If the ANOVA is significant, meaning the complex model shows a better fit than the reduced model, repeat steps above adding the subsequent regressor in the list for a newer complex model with one more component. If not, stop and model fitting is concluded. Once the best regressor order was determined, the stepwise regression continued with permutation tests to select optimal number of regressors: incrementally adding regressors in the optimal order until no more variance was captured (ANOVA p > 0.05).

#### Sum of F-stat statistic

To determine whether activity in a given frequency band encodes a specific regressor, we employed a sum-of-F-stat strategy. Briefly, a linear regression was calculated at each timepoint, resulting in a regression goodness-of-fit and significance value at each time point. Next, we examined the whole timecourse to determine the maximum number of consecutive regressions that resulted in a significant association (p < 0.05). We then calculated a summary statistic by summing the absolute value of the F-stat for each of those consecutive significant datapoints. This calculation has several benefits: it is agnostic to the exact timing of the activation, since it doesn’t impose any priors or temporal constraints on when it happens; it is sensitive to sustained activations, since those will result in several adjacent timepoints that cross the significant threshold; and it is sensitive to the strength of the activation, since greater correlations will result in higher F-stat values that add weight to the summary statistic. The resulting sum-of-F-stat therefore reflects the strength of the neural-behavioral association.

To obtain a significance estimation for the summary F-stat, we generate a Null sum-of-F-stat distribution by recalculating it after shuffling the behavioral labels 1000 times and carrying out the same calculation for each permutation. The resulting permutation p-value is the proportion of shuffled F-stats with a value higher than the observed sum-of-F-stat. If the activation is robust, this will result in a small p value since the majority of shuffled sum-of-F-stats will be well below the observed value.

### Decoding

#### Identifying optimal window of interest

To identify the optimal window of time before choice, we tested performance of a Euclidean distance classifier (ED, see below) on the first 50 ms before choice and then repeatedly increased the size of the window by 500 ms increments up to 2 s prior to choice. This was done separately for each subject, using all available electrodes for all six power bands and all available trials with leave-one-out (LOO) cross-validation. Classifier performance peaked at approximately 1 s prior to choice and for all subsequent analyses, we used a 50 ms resolution(17) and 1 s window prior to choice. Results were similar if we started 2 s before choice and added 50 ms windows to reach the time of choice (Supplementary Methods Figure 4). The resulting classification had modest performance, correctly classifying above chance in 18/20 patients (mean classification accuracy = 59.3±0.06%, p<0.001 vs bootstrapped performance).

#### Feature Selection

Choice active results (Fig. 3) indicated that choice-related information was distributed across all regions analyzed and was most likely to be carried by higher frequency bands (HFA, gamma, and possibly beta). We tested the contribution of beta to classification performance using linear dynamical systems (LDS) modeling and dynamic time warping (DTW) (see below) with gamma/HFA, with and without beta; classification results were nearly identical (R^2^ = 0.75) in both cases and not significantly different (p = 0.60) (Extended Data Table 22). Therefore, we selected gamma and HFA power bands in all regions for subsequent choice decoding.

#### Dimensionality Reduction

To reduce the dimensionality of data used for classification, we used two different dimensionality reduction techniques: principal component analysis (PCA) and linear dynamical systems modeling (LDS).

##### Principal component analysis (PCA)

To account for the variation in anatomical coverage and number of electrodes across patients, we performed PCA separately for each region in each patient. For each region, the original neural features were power in the gamma/HFA frequency bands for all electrodes. The original dataset for each patient thus contained a variable number of electrodes across a variable set of regions, and two power estimates (gamma and HFA). Through PCA, we reduced the number of features to the (varying) number of regions for each subject times the number of bands used (2, gamma and HFA). To avoid biasing the decoder based on whether the selected electrodes represented choice information, we included all electrodes in these analyses.

##### Linear dynamical system (LDS) model

As an alternative to PCA, we employed linear dynamical systems (LDS) modeling. LDS models allow characterization of the development of neural activity through time, and thus may be better suited for identifying how neural computations related to the on-going task unfold(83). As with PCA, LDS reduces the dimensionality of the data(84) but additionally uses an expectation-maximization likelihood function to solve for the parameters of the model. LDS have been widely used in the field of neuroscience to model the dynamics of neural activity. The underlying idea behind this approach is to capture the temporal dependencies and dynamics of neural signals using a linear combination of past activity and an input signal. LDS models have proven to be effective in characterizing the behavior of neural networks and predicting the response of neurons to external stimuli. In particular, it allows us to leverage the high temporal resolution of iEEG recordings to better understand the neural system dynamics. Due to its widespread usage, we can use well-studied methods specific to LDS, such as Lyapunov Exponents, in order to understand the neural dynamics across patients(35).

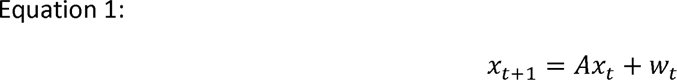

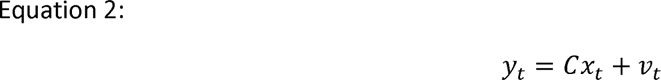

Equation 1 shows how the next state, *x_t_*_+1_, is calculated based on the previous state x_t_ and the state noise w_t_. Equation 2 calculates the output, y_t_, based on the state *x_t_* and output noise v_t_. Both A and C are learned using expectation maximization. The LDS seeks to determine a set of low-dimensional representations of neural activity that have maximal predictive power in their temporal trajectories. As above, the first PC for each region-powerband combination was used in LDS modeling. The appropriate dimensionality of the model (number of latent variables, LVs) was determined empirically by re-running the model for a range of dimensions (1–15) and examining the predictability of neural dynamics. When increasing the dimension size does not improve the predictability of the neural dynamics, the optimal dimensionality has been reached. The dimensionality value ranged from 8 to 16 across patients. In an effort to use a consistent number of dimensions for all patients, we selected a value of 7 LVs. Each dimension of the LDS model defines a latent variable that describes how the dynamics unfold and these LVs can be exploited to further probe the underlying dynamics of choice processing.

### Classifiers

One the dimensionality of the neural data was reduced using either PCA or LDS, we employed classification techniques to attempt to decode trial-by-trial patient choices from the neural data alone. Again, we employed two different strategies: a simple Euclidean Distance classifier and a Dynamic Time Warping approach.

#### Euclidean Distance Classifier (ED)

We used a simple Euclidean distance (ED) classifier as this method is computationally efficient and shown to be as effective as linear discriminate analysis (LDA) at classifying neural activity in response to different events(34). To estimate the decoding accuracy, we used a leave-one-out procedure. Briefly, we created separate neural trajectory templates for safe bets and gambles by averaging the dimensionality-reduced neural features (PCs or LVs, of size n ROIs x n frequencies (2) x n time bins) across trials of either type for each electrode, on a per patient basis. A single test trial of similar dimensions was left out from both the dimensionality reduction and this averaging procedure. After the templates were calculated, the overall Euclidean distance between this trial and the average safe bet/gamble templates was calculated. The predicted choice was then assigned to the closest template in Euclidean space. If the predicted choice matched the true, behaviorally expressed choice (either safe bet or gamble), the trial was correctly classified. This procedure was repeated for all trials in our sample. Performance was defined as the percentage of correctly classified trials, which is calculated on a patient-by-patient basis.

#### Dynamic Time Warping (DTW)

While an efficient classification method, the ED classifier’s fixed template matching and temporal comparison may not be the best approach for decoding complex neural processes. We attempted to improve on the ED classifier by using dynamic time-warping (DTW), a distance metric that is capable of accounting for variation in time of neural activation through flexible stretching or shrinking of the time dimension. Briefly, DTW seeks to minimize differences between two temporal sequences (in this case, the individual trial and each of the two templates) while allowing some flexibility in the matching. Specifically, instead of matching point-by-point, temporal warping that respects the sequence of points in each time series is allowed. DTW does this by using dynamic programming to calculate whether one or neither of the time series should be stretched in order to minimize the Euclidean distance between the two time series for each successive point in time, we used the Matlab implementation of dynamic time warping (`dtw`). We hypothesized this additional temporal flexibility would result in classification improvement if the timing of neural activity supporting patient choices varies from trial to trial, but the underlying computations are the same. For DTW we used the same leave-one-out procedure as the ED-based classifier with the average for both safe and gamble trials separately warped with the left-out trial before calculating the ED.

### Bootstrapping

To identify bias in the neural data that may account for performance above the expected 50% chance performance, we randomly swapped the labels (gamble or safe bet) for each trial and assessed performance on the PCA+LDS+DTW model repeatedly, 1000 times (e.g. bootstrap performance). The distribution of performance provides an empirical assessment of chance performance for each subject. This is especially important here because in this task, subjects are more likely to select a gamble than a safe bet (see Results) which could bias classifier performance. The bootstrap performance across subjects was almost exactly 50%, as expected (50.1±0.002%), whereas the actual performance was significantly better than chance (74.3±3.4%, paired t-test, p < .01). In addition, we calculated decoder performance using balanced accuracy. Balanced accuracy is calculated by calculating the accuracy separately for each class (gamble and safe bet), and averaging them.

#### Latent variable trajectories and manifolds

We examined the latent variables (LVs) from the LDS model in order to probe the underlying choice dynamics. For plotting purposes, three of seven LVs from one patient were selected that maximized differences (ED) between gamble and safe bet trials. The patient with the best decoder performance was chosen (p06, 77.0%, see Extended Methods Table 16). The selected LVs for all gamble trials and safe bet trials were then averaged and graphed, illustrating the differences in LVs between safe bets and gamble trials (Fig. 4C). A manifold was defined by an ellipsoid that contained the average gamble and safe bet trajectories for the selected patient (min and max for each LV dimension plus an additional 70% (LV_1_) or 60% (LV_2_, LV_3_).

Whereas the classification results can be compared across subjects, the visualization of the statespace trajectories with time are not comparable across patients. To address this issue, we estimated and compared Lyapunov exponents, which measure the rate at which two initially similar multi-dimensional trajectories diverge from each other as time progresses(35) with positive values indicating that the trajectories are getting farther apart over time, e.g. more chaotic, and negative values indicating the trajectories are getting closer together over time, or converging on an attractor. To accomplish this, for each of the 7-dimensional trajectories, for each trial, we calculated the rate of divergence between the original LV and a perturbed version of the LV to estimate the Lyapunov exponent over time separately for each subject. The temporal evolution of Lyapunov exponents was averaged across subjects to evaluate whether, on average across subjects, the Lyapunov exponent declined towards negative values, indicating an attractor.

### Contribution of brain areas/subcircuits to decoding accuracy

To examine the contribution of individual brain areas and subcircuits to decoding, we calculated model decoding accuracy after progressively adding individual regions/subcircuits. We started by calculating model decoding accuracy as above using only the region with the worst individual classifier performance, and iteratively added regions in order of increasing performance until all regions were included, separately for each subject we sought to examine the contribution. To examine the contribution of subcircuits, we performed the same analysis using subcircuits (prefrontal (OFC/LPFC/CC), frontoparietal (PrG/PoG/PC), and limbic (Amy/HC/Ins)) instead of regions.

