## Supplemental Figures and Tables for "Multi-areal neural dynamics encode human decision making"

List of Extended Data figures and tables

### Extended Data figures & tables

#### Extended Data Figure 1: Individual patient gambling behavior.

Light grey datapoints show individual trial choices (light grey; 1=gamble, 0=safe bet). Black points show averaged gamble proportion for all trials in 10% win probability bins. All patients gamble more often when win probability is higher. Red line indicates sigmoidal logistic fit (all p<0.01), indicating that patients understood the task and played to maximize their winnings. Patients were excluded if their behavioral data could not be fit with a sigmoidal curve.

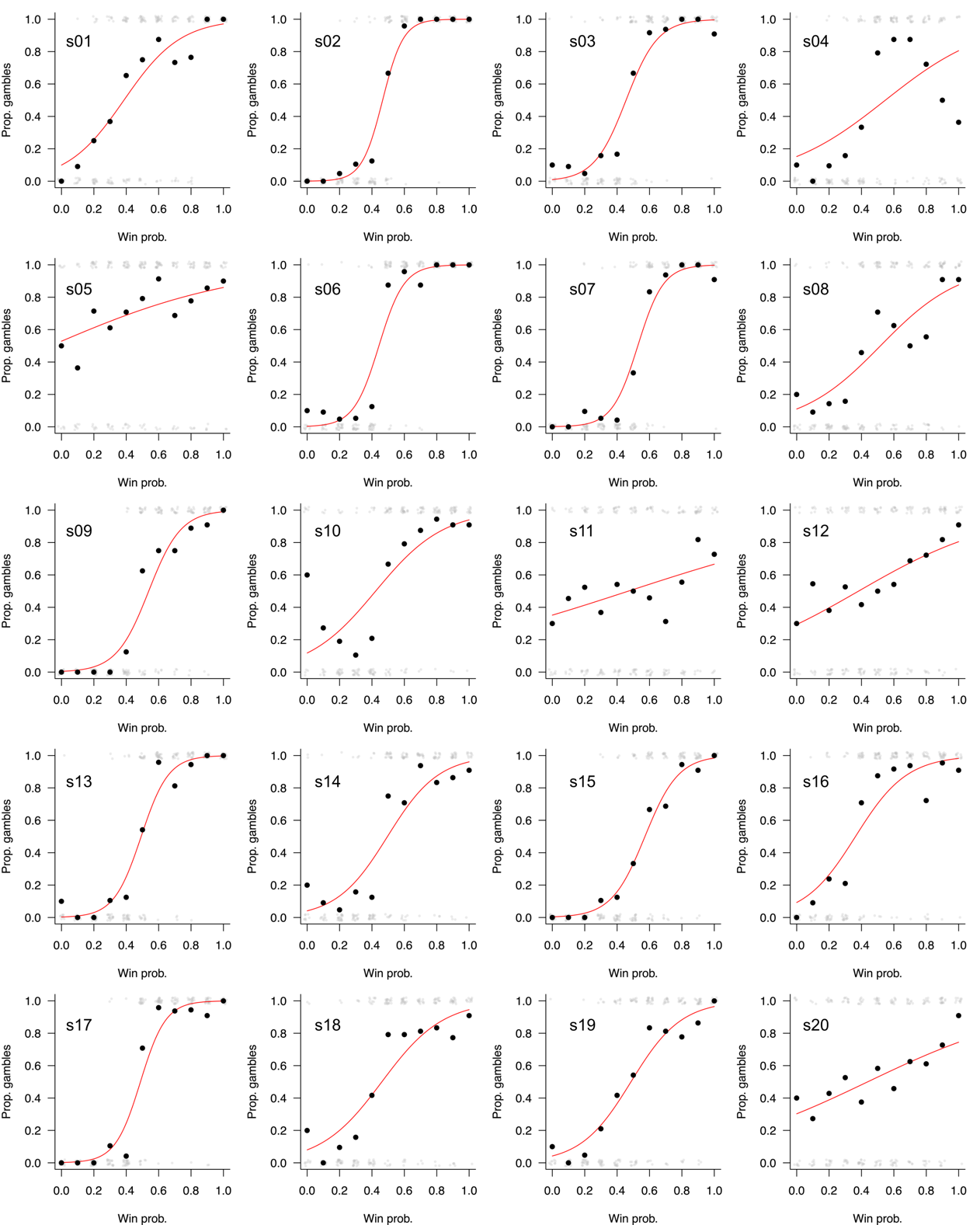

#### Extended Data Figure 2: Anatomical coverage for individual patients

Anatomical coverage for individual patients (p01-p20). Patients p01 through p09 underwent electrocorticography (ECoG) implantation, whereas patients p10-p20 underwent stereotactic EEG (sEEG) implantation. Areas of interest with 2+ electrodes were included for all patients. See Table P1 for details on electrode count by area and patient.

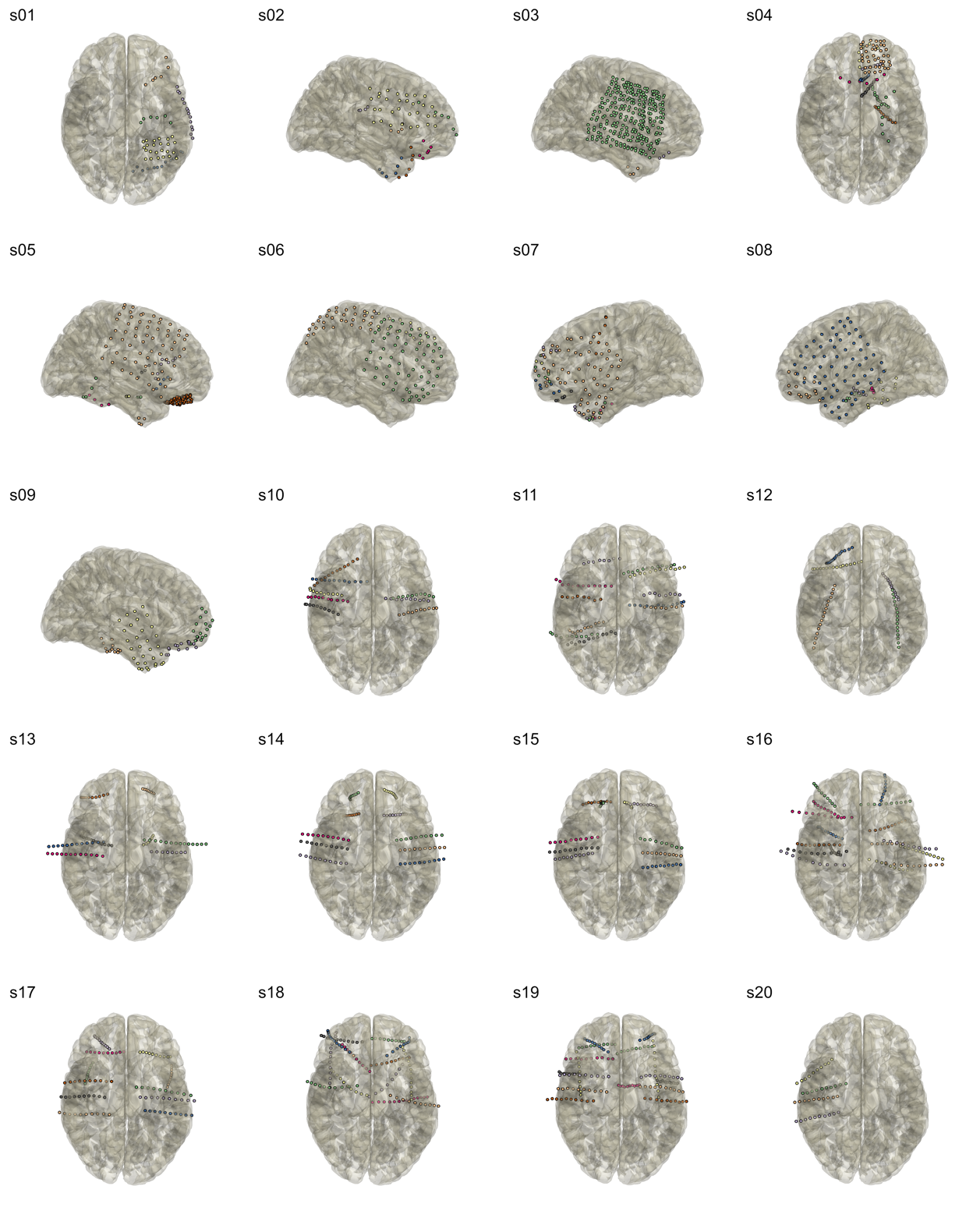

#### Extended Data Table 1: Count of electrodes per patient and region

Table shows the number of electrodes in each anatomical regions of interest for each patient in our final sample (n=20), as well as the total count of electrodes per ROI across all patients and the number of patients with coverage for each ROI.

| **Patient** | **LPFC** | **OFC** | **Cingulate** | **Amygdala** | **Hippocampus** | **Insula** | **Precentral Gyrus** | **Postcentral Gyrus** | **Parietal** | **Total electrode number** |
| --- | --- | --- | --- | --- | --- | --- | --- | --- | --- | --- |
| p01 | 0 | 1 | 0 | 0 | 1 | 0 | 0 | 0 | 13 | 15 |
| p02 | 24 | 5 | 0 | 0 | 0 | 0 | 7 | 8 | 2 | 46 |
| p03 | 55 | 0 | 0 | 0 | 0 | 0 | 47 | 36 | 22 | 160 |
| p04 | 7 | 49 | 6 | 1 | 7 | 0 | 0 | 0 | 0 | 70 |
| p05 | 30 | 61 | 3 | 0 | 0 | 5 | 14 | 8 | 5 | 126 |
| p06 | 29 | 7 | 0 | 0 | 0 | 0 | 10 | 15 | 24 | 85 |
| p07 | 41 | 11 | 0 | 0 | 0 | 0 | 11 | 3 | 2 | 68 |
| p08 | 26 | 10 | 2 | 1 | 8 | 0 | 9 | 6 | 4 | 66 |
| p09 | 15 | 14 | 0 | 0 | 0 | 0 | 1 | 2 | 0 | 32 |
| p10 | 8 | 3 | 2 | 6 | 6 | 0 | 1 | 0 | 0 | 26 |
| p11 | 11 | 0 | 11 | 4 | 3 | 3 | 2 | 6 | 6 | 46 |
| p12 | 15 | 5 | 4 | 5 | 9 | 5 | 0 | 0 | 0 | 43 |
| p13 | 9 | 3 | 1 | 4 | 0 | 0 | 0 | 0 | 0 | 17 |
| p14 | 15 | 4 | 4 | 1 | 4 | 0 | 0 | 0 | 0 | 28 |
| p15 | 11 | 3 | 8 | 5 | 6 | 0 | 0 | 0 | 0 | 33 |
| p16 | 12 | 4 | 3 | 2 | 10 | 1 | 3 | 0 | 0 | 35 |
| p17 | 11 | 2 | 8 | 2 | 7 | 6 | 0 | 0 | 0 | 36 |
| p18 | 29 | 6 | 18 | 0 | 1 | 7 | 3 | 4 | 0 | 68 |
| p19 | 38 | 4 | 14 | 1 | 2 | 14 | 0 | 0 | 0 | 73 |
| p20 | 5 | 1 | 0 | 0 | 1 | 5 | 0 | 0 | 0 | 12 |
| **# elecs per ROI** | **391** | **193** | **84** | **32** | **65** | **46** | **108** | **88** | **78** | **1085** |
| **# patients with ROI coverage** | **19** | **18** | **13** | **11** | **13** | **8** | **11** | **9** | **8** | **110** |

#### Extended Data Table 2: Reaction times

Reaction times means and standard deviation (seconds) for each patient. All completed trials with the exception of one outlier trial (p17, rt= 193.5 secs) are included regardless of whether the trial was excluded from analysis due to artifact or noise (3287 trials total). Overall mean reaction time = 1.00 seconds, stdev = 0.017.

| **Patient** | **Mean** | **stdev** | **N trials** |
| --- | --- | --- | --- |
| p01 | 1.21 | 0.295 | 180 |
| p02 | 1.05 | 0.248 | 188 |
| p03 | 0.88 | 0.259 | 177 |
| p04 | 0.84 | 0.147 | 194 |
| p05 | 0.87 | 0.306 | 176 |
| p06 | 1.04 | 0.307 | 187 |
| p07 | 1.25 | 0.482 | 181 |
| p08 | 1.71 | 0.514 | 196 |
| p09 | 1.35 | 0.410 | 200 |
| p10 | 1.32 | 0.390 | 195 |
| p11 | 1.58 | 0.477 | 74 |
| p12 | 2.30 | 0.742 | 38 |
| p13 | 1.35 | 0.750 | 200 |
| p14 | 1.16 | 0.407 | 172 |
| p15 | 2.02 | 0.784 | 172 |
| p16 | 0.80 | 0.259 | 200 |
| p17 | 1.33 | 1.166 | 53 |
| p18 | 1.89 | 1.214 | 168 |
| p19 | 0.80 | 0.212 | 197 |
| p20 | 2.27 | 1.039 | 139 |
| Overall | 1.00 | 0.017 | 3287 |

#### Extended Data Figure 3: Distribution of reaction times

Histogram shows distribution of reaction times over all trials as above completed for all patients (3287 trials, n=20 patients).

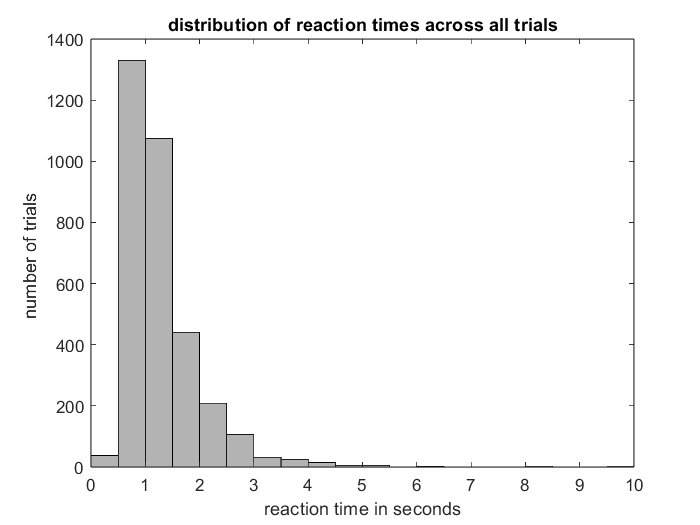

#### Extended Data Figure 4: Temporal window analysis

Plot shows average performance (proportion correct classification) across subjects of an increasing window size towards choice (0 s, light green), or start from choice backwards to -2 s (dark green). Classifier performance peaked at approximately 1 s prior to choice and for all subsequent analyses, we used a 50 ms resolution and 1 s window prior to choice. Results were similar if we started 2 s before choice and added 50 ms windows to reach the time of choice.

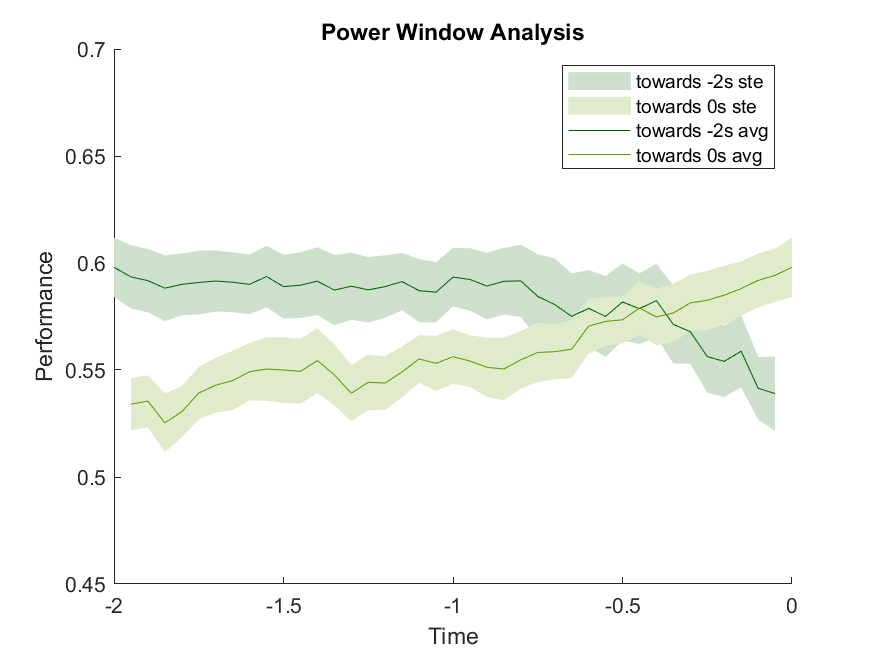

#### Extended Data Table 3: Counts and percentages of task active electrodes per region

Table shows the number of electrodes in each anatomical regions of interest: total number and number/proportion showing significant power modulation in any frequency band during deliberation, per ROI.

|  | Total # Electrodes | # task active electrodes | Proportion of task active electrodes |
| --- | --- | --- | --- |
| Amygdala | 27 | 19 | 70.4% |
| Hippocampus | 56 | 38 | 67.9% |
| Insula | 41 | 32 | 78.0% |
| Cingulate | 80 | 60 | 75.0% |
| LPFC | 376 | 284 | 75.5% |
| OFC | 188 | 126 | 67.0% |
| Parietal | 78 | 75 | 96.2% |
| PostcentralG | 88 | 74 | 84.1% |
| PrecentralG | 108 | 92 | 85.2% |
| Grand Total | 1042 | 800 | 76.8% |

#### Extended Data Table 4: Counts and proportions of task active electrodes for each patient

Table shows the number of electrodes in each anatomical regions of interest: total number and number/proportion showing significant power modulation in any frequency band during deliberation, per patient.

| Patient | Total # Electrodes | # task active electrodes | Proportion of task active electrodes |
| --- | --- | --- | --- |
| p01 | 15 | 15 | 1.000 |
| p02 | 46 | 41 | 0.891 |
| p03 | 160 | 133 | 0.831 |
| p04 | 70 | 51 | 0.729 |
| p05 | 126 | 73 | 0.579 |
| p06 | 85 | 77 | 0.906 |
| p07 | 68 | 22 | 0.324 |
| p08 | 66 | 58 | 0.879 |
| p09 | 32 | 32 | 1.000 |
| p10 | 26 | 23 | 0.885 |
| p11 | 46 | 41 | 0.891 |
| p13 | 17 | 12 | 0.706 |
| p14 | 28 | 23 | 0.821 |
| p15 | 33 | 22 | 0.667 |
| p16 | 35 | 30 | 0.857 |
| p17 | 36 | 19 | 0.528 |
| p18 | 68 | 57 | 0.838 |
| p19 | 73 | 71 | 0.973 |
| p20 | 12 | 0 | 0.000 |
| Mean | 54.8 | 42.1 | 0.753 |

#### Extended Data Table 5: Counts of task active electrodes for each region and number of active powers

Table shows the number of electrodes showing significant power modulation in none, one, or several frequency bands. For each ROI, the number of electrodes showing activation in 0-6 frequency bands is indicated. On average there were 2.09±1.7 active bands per electrode. 242 electrodes (23.2%) showed no power modulation in any frequency band, whereas active electrodes (with one or more active power bands) have on average 2.72 active power bands.

| Regions | Number of task-active frequency bands | | | | | | | Mean |
| --- | --- | --- | --- | --- | --- | --- | --- | --- |
|  | 0 | 1 | 2 | 3 | 4 | 5 | 6 |  |
| Amygdala | 8 | 6 | 4 | 5 | 1 | 2 | 1 | 1.81 |
| Hippocampus | 18 | 11 | 17 | 7 | 2 | 1 | 0 | 1.41 |
| Insula | 9 | 17 | 8 | 2 | 2 | 3 | 0 | 1.51 |
| OFC | 62 | 34 | 32 | 18 | 23 | 18 | 1 | 1.81 |
| Cingulate | 20 | 17 | 14 | 17 | 8 | 3 | 1 | 1.86 |
| LPFC | 92 | 71 | 71 | 64 | 41 | 31 | 6 | 2.02 |
| PrecentralG | 16 | 21 | 28 | 21 | 13 | 5 | 4 | 2.23 |
| PostcentralG | 14 | 15 | 12 | 24 | 7 | 10 | 6 | 2.56 |
| Parietal | 3 | 3 | 14 | 24 | 8 | 17 | 9 | 3.51 |
| Sum | 242 | 195 | 200 | 182 | 105 | 90 | 28 | 2.09 |

#### Extended Data Table 6: Proportion of task active electrodes per frequency band (patient means & SEs)

Table shows the mean proportion of task-active electrodes showing significant power modulation in each frequency band across patients (Figure 2b. The frequency with the highest proportion of task-active electrodes was HFA (39.0%), whereas delta showed the lowest (28.7%).

| Frequency band | Mean across Patients | Standard deviation | Standard error |
| --- | --- | --- | --- |
| Delta | 28.7% | 0.252 | 0.058 |
| Theta | 36.4% | 0.249 | 0.057 |
| Alpha | 32.4% | 0.186 | 0.043 |
| Beta | 36.5% | 0.256 | 0.059 |
| Gamma | 32.6% | 0.226 | 0.052 |
| HFA | 39.0% | 0.226 | 0.052 |

#### Extended Data Table 7: Proportion of task active electrodes per frequency band and region

Proportion of task-active electrodes per region and frequency band. The frequency with the highest proportion of task-active electrodes was beta (41.2%), whereas delta showed the lowest (25.8%). Numbers are out of the total number of electrodes per regions, whereas numbers in parentheses indicate the proportion out of the number of task-active electrodes per region.

| Regions | Delta | Theta | Alpha | Beta | Gamma | HFA |
| --- | --- | --- | --- | --- | --- | --- |
| Amygdala | 0.111 (0.158) | 0.333 (0.474) | 0.333 (0.474) | 0.259 (0.368) | 0.296 (0.421) | 0.481 (0.684) |
| Hippocampus | 0.321 (0.474) | 0.304 (0.447) | 0.25 (0.368) | 0.286 (0.421) | 0.054 (0.079) | 0.196 (0.289) |
| Insula | 0.146 (0.188) | 0.366 (0.469) | 0.195 (0.250) | 0.415 (0.531) | 0.171 (0.219) | 0.220 (0.281) |
| OFC | 0.277 (0.413) | 0.309 (0.460) | 0.287 (0.429) | 0.106 (0.159) | 0.399 (0.595) | 0.431 (0.643) |
| Cingulate | 0.238 (0.317) | 0.413 (0.550) | 0.338 (0.450) | 0.400 (0.533) | 0.200 (0.267) | 0.275 (0.367) |
| LPFC | 0.290 (0.384) | 0.327 (0.433) | 0.250 (0.331) | 0.356 (0.472) | 0.351 (0.465) | 0.447 (0.592) |
| PrecentralG | 0.130 (0.152) | 0.231 (0.272) | 0.343 (0.402) | 0.704 (0.826) | 0.509 (0.598) | 0.315 (0.370) |
| PostcentralG | 0.239 (0.284) | 0.295 (0.351) | 0.557 (0.662) | 0.648 (0.770) | 0.432 (0.514) | 0.386 (0.459) |
| Parietal | 0.346 (0.360) | 0.526 (0.547) | 0.679 (0.707) | 0.897 (0.933) | 0.603 (0.627) | 0.462 (0.480) |
| Mean | 0.258 (0.336) | 0.333 (0.434) | 0.331 (0.431) | 0.412 (0.536) | 0.366 (0.476) | 0.392 (0.510) |

#### Extended Data Table 8: Proportion of task active electrodes per region

Mean proportion task active electrodes across patients (any frequency band) by ROI (Figure 2c).

| ROI | Mean across patients (any frequency band) | Standard deviation | Standard error |
| --- | --- | --- | --- |
| Amygdala | 63.0% | 0.397 | 0.126 |
| Hippocampus | 69.9% | 0.358 | 0.103 |
| Insula | 79.3% | 0.358 | 0.135 |
| OFC | 77.4% | 0.289 | 0.070 |
| Cingulate | 77.8% | 0.323 | 0.093 |
| LPFC | 77.1% | 0.277 | 0.065 |
| PrG/Motor | 86.2% | 0.198 | 0.060 |
| PoG/Somato | 81.0% | 0.231 | 0.077 |
| Parietal | 81.3% | 0.372 | 0.132 |

#### Extended Data Figure 5. Proportion of task responsive electrodes: increased vs decreased.

Proportion of task responsive electrodes, defined as electrodes that show a significant power modulation during deliberation compared to baseline. These data correspond to that in Fig. 2D bar plot but organized to show patterns across frequency bands. Data also correspond to scatterplots in Fig 2F,G. All error bars indicate standard error of the mean.

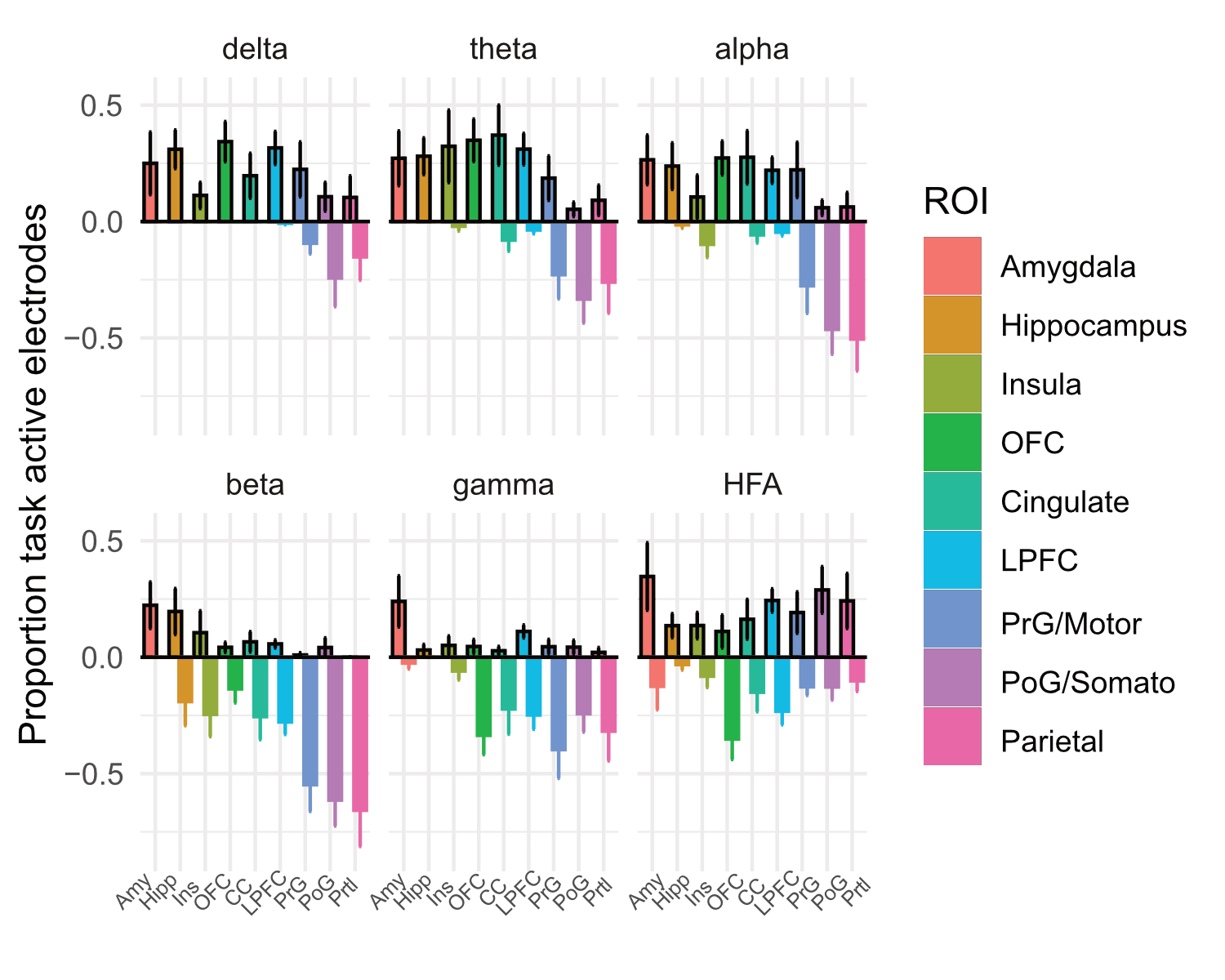

#### Extended Data Table 9: Count of electrodes that increased power, decreased power, both (in separate bands), or neither, per region

Table shows the number of electrodes that showed power modulation, separated by whether the modulation was only power increases, decreases, both or none. Results reflect modulation in any frequency band, such that decrease in one frequency and decreases could appear simultaneously in different frequency bands.

|  | **Proportion of electrodes** | | | | |
| --- | --- | --- | --- | --- | --- |
| **Region** | **Power decreases only** | **Power increases only** | **Power increases AND decreases** | **No power modulation** | **Total** |
| Amygdala | 3.7% | 59.3% | 7.4% | 29.6% | 100% |
| Cingulate | 23.8% | 25.0% | 26.3% | 25.0% | 100% |
| Hippocampus | 7.1% | 44.6% | 16.1% | 32.1% | 100% |
| Insula | 24.4% | 29.3% | 24.4% | 22.0% | 100% |
| LPFC | 19.4% | 23.9% | 32.2% | 24.5% | 100% |
| OFC | 19.1% | 18.6% | 29.3% | 33.0% | 100% |
| Parietal | 59.0% | 2.6% | 34.6% | 3.8% | 100% |
| PostcentralG | 34.1% | 12.5% | 37.5% | 15.9% | 100% |
| PrecentralG | 50.9% | 9.3% | 25.0% | 14.8% | 100% |
| **Grand Total** | **26.3%** | **21.2%** | **29.3%** | **23.2%** | **100%** |

#### Extended Data Table 10: Counts and percentages of electrodes that showed power increases/decreases per frequency band

Electrode counts and % of total electrodes (n=1042) by frequency band that decreased or increased power relative to baseline.

|  | **Count of Elecs that** | | **% of Total Elecs that** | |
| --- | --- | --- | --- | --- |
| **Row Labels** | **Decreased Power** | **Increased Power** | **Decreased Power** | **Increased Power** |
| Delta | 53 | 216 | 5.1% | 20.7% |
| Theta | 97 | 250 | 9.3% | 24.0% |
| Alpha | 156 | 189 | 15.0% | 18.1% |
| Beta | 367 | 62 | 35.2% | 6.0% |
| Gamma | 306 | 75 | 29.4% | 7.2% |
| HFA | 208 | 200 | 20.0% | 19.2% |

#### Extended Data Table 11: Clustering Task-Active Electrodes

Alignment results of k-nearest neighbor clustering UMAP projections with the three clusters (columns) with the three proposed groups of regions (rows): 46.8% of the points in cluster one belonged to frontoparietal regions (PrG, PoG, PC), 45.8% of cluster 2 datapoints belonged to limbic regions (Amy, HC, Ins), and 53.6% of cluster 3 datapoints belonged to prefrontal regions (OFC, LPFC, CC).

|  | Cluster 1 | Cluster 2 | Cluster 3 |
| --- | --- | --- | --- |
| Prefrontal | 0.213 | 0.197 | **0.536** |
| Limbic | 0.319 | **0.458** | 0.214 |
| Fronto-Parietal | **0.468** | 0.345 | 0.250 |

#### Extended Data Table 12: Counts & percentages of choice-active electrodes overall

Tables shows counts of electrodes that show no choice activity (0 frequency bands) and count of electrodes with significant choice-activity in 1 or more frequency bands. Right columns chow percentage of choice-activeelectrodes per region, and mean number of active frequency bands per region, including and excluding electrodes with no choice-related activity.  On average across all electrodes, including those that were not active, 42.7% were choice-active and there were 0.55 active bands per electrode. Active electrodes (with one or more active power bands) have on average 1.29 active power bands (i.e., excluding electrodes with 0 active powers); 597/1042 = 57.3% of electrodes were not active in any power band.

|  | Total # electrodes | Number of Choice-Active Frequency Bands (no electrodes had 5 or 6 active bands) | | | | | % of Choice-Active Electrodes (≥ 1 freq band) | Mean # Choice-Active frequency bands (including 0) | Mean # Choice-Active bands (excluding 0) |
| --- | --- | --- | --- | --- | --- | --- | --- | --- | --- |
|  |  | 0 | 1 | 2 | 3 | 4 |  |  |  |
| Amygdala | 27 | 18 | 8 | 1 | 0 | 0 | 33.3% | 0.37 | 1.11 |
| Hippocampus | 56 | 38 | 14 | 3 | 1 | 0 | 32.1% | 0.41 | 1.28 |
| Insula | 41 | 27 | 11 | 2 | 1 | 0 | 34.1% | 0.44 | 1.29 |
| OFC | 188 | 103 | 69 | 12 | 4 | 0 | 45.2% | 0.56 | 1.24 |
| Cingulate | 80 | 51 | 23 | 6 | 0 | 0 | 36.3% | 0.44 | 1.21 |
| LPFC | 376 | 229 | 112 | 33 | 1 | 1 | 39.1% | 0.49 | 1.26 |
| PrecentralG | 108 | 54 | 43 | 9 | 2 | 0 | 50.0% | 0.62 | 1.24 |
| PostcentralG | 88 | 42 | 30 | 15 | 1 | 0 | 52.3% | 0.72 | 1.37 |
| Parietal | 78 | 35 | 22 | 16 | 5 | 0 | 55.1% | 0.88 | 1.6 |
| Sum \| [Mean] | 1042 | 597 | 332 | 97 | 15 | 1 | [42.7%] | [0.55] | [1.29] |

#### Extended Data Table 13: Counts & proportions of choice-active electrodes per patient

Tables show total electrodes, count, and proportion of choice-active electrodes per patient.

Across patients, the mean proportion choice-encoding electrodes across patients was 0.375 (SE = 0.028; Fig 3a).

| Patient | Total Electrodes | Number of Choice-active electrodes | Proportion Choice-active Electrodes |
| --- | --- | --- | --- |
| p01 | 15 | 5 | 0.333 |
| p02 | 46 | 19 | 0.413 |
| p03 | 160 | 82 | 0.513 |
| p04 | 70 | 25 | 0.357 |
| p05 | 126 | 82 | 0.651 |
| p06 | 85 | 34 | 0.400 |
| p07 | 68 | 31 | 0.456 |
| p08 | 66 | 28 | 0.424 |
| p09 | 32 | 7 | 0.219 |
| p10 | 26 | 10 | 0.385 |
| p11 | 46 | 17 | 0.370 |
| p13 | 17 | 8 | 0.471 |
| p14 | 28 | 11 | 0.393 |
| p15 | 33 | 8 | 0.242 |
| p16 | 35 | 13 | 0.371 |
| p17 | 36 | 11 | 0.306 |
| p18 | 68 | 18 | 0.265 |
| p19 | 73 | 35 | 0.479 |
| p20 | 12 | 1 | 0.083 |

#### Extended Data Table 14: Proportion of choice-active electrodes per power band (patient means & SEs)

Data Table corresponds to Figure 3C: mean proportion choice-active electrodes +/- SE per frequency band. Greatest to least: HFA (19.7%), gamma (13.3%), beta (6.05%), alpha (3.24%), theta (2.14%), and delta (2.76%). Significantly greater proportion of electrodes represented choice in the higher frequencies than lower frequencies (p < 0.05E-7, t-test, 2-tailed t-test, 2-sample unequal variances).

| Frequency band | Mean across patients | SDs | SEs |
| --- | --- | --- | --- |
| Delta | 2.76% | 4.51% | 1.03% |
| Theta | 2.14% | 2.48% | 0.57% |
| Alpha | 3.24% | 4.37% | 1.00% |
| Beta | 6.05% | 5.69% | 1.31% |
| Gamma | 13.3% | 6.29% | 1.44% |
| HFA | 19.7% | 9.27% | 2.13% |

#### Extended Data Table 15: Proportion of choice-active electrodes per region (patient means & SEs)

Data Table shows per region mean proportion of choice-active electrodes, standard deviations, and standard errors, across subjects. Data corresponds to Figure 3b.

| ROI | Means | SDs | SEs |
| --- | --- | --- | --- |
| Amygdala | 0.277 | 0.353 | 0.112 |
| Hippocampus | 0.221 | 0.221 | 0.064 |
| Insula | 0.342 | 0.321 | 0.121 |
| OFC | 0.424 | 0.321 | 0.078 |
| Cingulate | 0.381 | 0.305 | 0.088 |
| LPFC | 0.357 | 0.163 | 0.039 |
| PrecentralG | 0.477 | 0.361 | 0.109 |
| PostcentralG | 0.410 | 0.325 | 0.108 |
| Parietal | 0.442 | 0.318 | 0.112 |

#### Extended Data Table 16: Behavioral Regressors

| **Regressor** | **Formula** | **Description** |
| --- | --- | --- |
| Win probability | $p$ | Probability of gamble resulting in a win (e.g., 70% for a shown 3), regardless of choice |
| Risk | $\boldsymbol{E}\{\left( \boldsymbol{V}-\boldsymbol{E}\left( \boldsymbol{V} \right) \right)^{2}\}$ | Risk (level of uncertainty) associated with the gamble option. Risk is maximal when win probability is 0.5, and 0 when win probability is 0 or 1. |
| Risky side | $\left\{ \begin{matrix} 1 & gamble on right \\ 0 & gamble on left \end{matrix} \right.$ | Side of the screen that gamble option was presented on a given trial: 1 if on the right side, 0 if on the left. |
| Choice side | $\left\{ \begin{matrix} 1 & chose right \\ 0 & chose left \end{matrix} \right.$ | Side of screen that patient chose (or button selected) on a given trial: 1 if right side, 0 if left. |
| Choice | $\left\{ \begin{matrix} 1 & chose gamble \\ 0 & chose safe bet \end{matrix} \right.$ | Boolean indicator for gamble choice: 1 if patient chose to gamble, else 0 for safebet. |

#### Extended Data Figure 6: Cross correlations among regressors of interest pooled across patients (n=20)

Cross correlation matrix among six behavioral regressors of interest studied. All off diagonal R^2^ values were less than or equal to 0.01, except for win probability and choice (R^2^ = 0.34), indicating low cross-regressor collinearity.

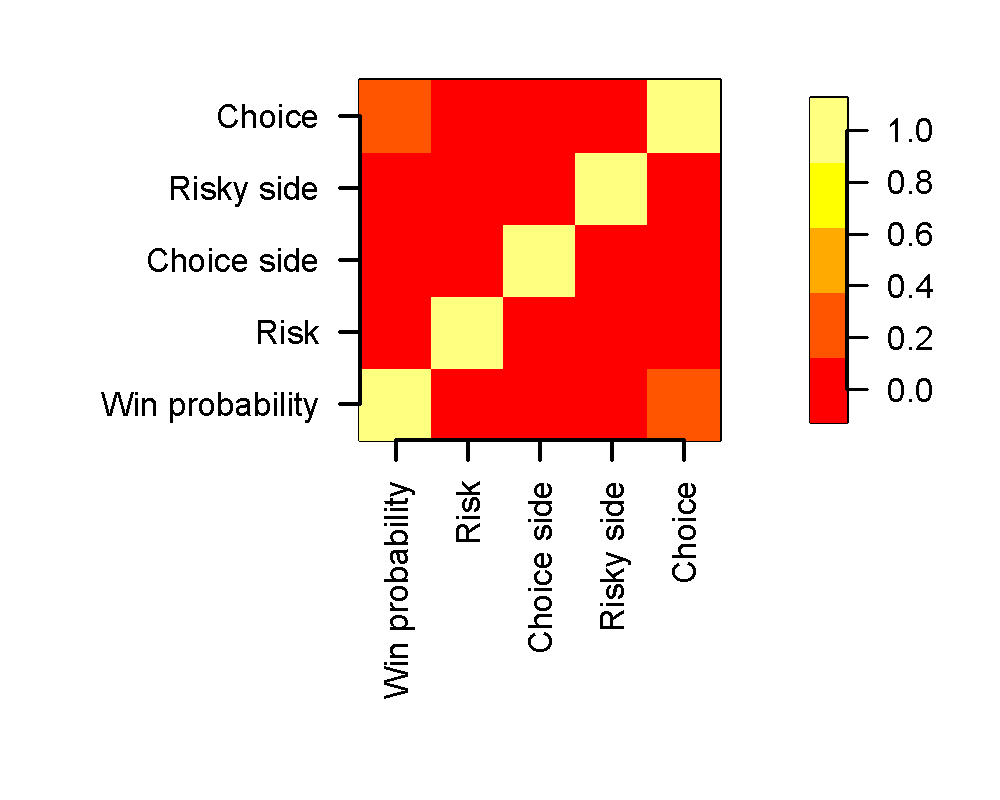

#### Extended Data Table 17: Electrode Counts per patient and region of interest included in Stepwise Regression Analyses

| Patient | **Hipp** | **OFC** | **PL** | **LPFC** | **PrG** | **PoG** | **Amyg** | **CG** | **INS** | **Total** |
| --- | --- | --- | --- | --- | --- | --- | --- | --- | --- | --- |
| p01 | 1 | 1 | 13 | *NA* | *NA* | *NA* | *NA* | *NA* | *NA* | 15 |
| p02 | *NA* | 5 | 2 | 24 | 7 | 8 | *NA* | *NA* | *NA* | 46 |
| p03 | *NA* | *NA* | 22 | 55 | 47 | 36 | *NA* | *NA* | *NA* | 160 |
| p04 | 7 | 49 | *NA* | 7 | *NA* | *NA* | 1 | 6 | *NA* | 70 |
| p05 | *NA* | 61 | 5 | 30 | 14 | 8 | *NA* | 3 | 5 | 126 |
| p06 | *NA* | 7 | 24 | 29 | 10 | 15 | *NA* | *NA* | *NA* | 85 |
| p07 | *NA* | 11 | 2 | 41 | 11 | 3 | *NA* | *NA* | *NA* | 68 |
| p08 | 8 | 10 | 4 | 26 | 9 | 6 | 1 | 2 | *NA* | 66 |
| p09 | *NA* | 14 | *NA* | 15 | 1 | 2 | *NA* | *NA* | *NA* | 32 |
| p10 | 6 | 3 | *NA* | 8 | 1 | *NA* | 6 | 2 | *NA* | 26 |
| p13 | *NA* | 3 | *NA* | 9 | *NA* | *NA* | 4 | 1 | *NA* | 17 |
| p16 | 10 | 4 | *NA* | 12 | 3 | *NA* | 2 | 3 | 1 | 35 |
| p18 | 1 | 6 | *NA* | 29 | 3 | 4 | *NA* | 18 | 7 | 68 |
| p19 | 2 | 4 | *NA* | 38 | *NA* | *NA* | 1 | 14 | 14 | 73 |
| **Total** | 35 | 178 | 72 | 323 | 106 | 82 | 15 | 49 | 27 | **887** |

#### Extended Data Table 18: % encoding electrodes per region and regressor

| **ROI** | **Win probability** | **Risk** | **Risky side** | **Choice side** | **Choice** |
| --- | --- | --- | --- | --- | --- |
| Hippocampus | 34.3% | 20.0% | 20.0% | 17.1% | 42.9% |
| Amygdala | 26.7% | 26.7% | 40.0% | 33.3% | 60.0% |
| Insula | 18.5% | 29.6% | 48.1% | 29.6% | 55.6% |
| Cingulate | 22.4% | 38.8% | 40.8% | 36.7% | 34.7% |
| OFC | 31.5% | 47.2% | 27.0% | 27.0% | 50.6% |
| LPFC | 36.2% | 36.2% | 24.8% | 23.8% | 43.0% |
| Parietal | 40.3% | 30.6% | 36.1% | 40.3% | 55.6% |
| PrecentralG | 38.7% | 25.5% | 29.2% | 37.7% | 44.3% |
| PostcentralG | 28.0% | 25.6% | 28.0% | 51.2% | 45.1% |

#### Extended Data Table 19: Decoder performance per patient, excluding trials with no uncertainty

Performance (% accurately classified trials) for each decoder model for each patient. Trials with no uncertainty (i.e., gamble win probabilities of 0% and 100%) are excluded. Data Table corresponds to Figure 4a. Abbreviations: latent variables (LV), principal components (PC), Euclidean distance (ED), dynamic time warping (DTW).

| Patient | LV + DTW | LV + ED | PC + DTW | PC + ED | Best |
| --- | --- | --- | --- | --- | --- |
| p01 | 65.7% | 68.1% | 63.8% | 60.3% | 68.1% |
| p02 | 76.5% | 61.9% | 57.9% | 67.4% | 76.5% |
| p03 | 72.5% | 62.9% | 61.9% | 70.8% | 72.5% |
| p04 | 76.1% | 56.9% | 63.6% | 57.5% | 76.1% |
| p05 | 77.1% | 62.8% | 74.0% | 69.3% | 77.1% |
| p06 | 79.7% | 77.7% | 77.1% | 68.8% | 79.7% |
| p07 | 76.7% | 64.8% | 54.0% | 55.9% | 76.7% |
| p08 | 71.9% | 65.7% | 67.3% | 61.8% | 71.9% |
| p09 | 77.4% | 78.5% | 68.5% | 58.0% | 78.5% |
| p10 | 79.8% | 71.3% | 60.7% | 53.5% | 79.8% |
| p11 | 71.5% | 66.1% | 62.8% | 65.9% | 71.5% |
| p12 | 75.2% | 73.4% | 56.9% | 64.1% | 75.2% |
| p13 | 73.6% | 72.1% | 59.0% | 50.6% | 73.6% |
| p14 | 70.5% | 63.1% | 61.8% | 57.4% | 70.5% |
| p15 | 76.6% | 78.3% | 54.7% | 54.2% | 78.3% |
| p16 | 72.3% | 71.1% | 64.1% | 65.2% | 72.3% |
| p17 | 76.1% | 75.3% | 55.8% | 40.5% | 76.1% |
| p18 | 72.4% | 70.1% | 63.6% | 60.1% | 72.4% |
| p19 | 73.1% | 71.1% | 70.2% | 63.8% | 73.1% |
| p20 | 70.8% | 71.5% | 52.9% | 47.7% | 71.5% |
| Mean | 74.3% | 69.1% | 62.5% | 59.6% | 74.6% |
| SD | 3.36% | 5.86% | 6.36% | 7.60% | 3.19% |

#### Extended Data Table 20: Decoder performance across Win Probabilities

Performance (% accurately classified trials) for LV+DTW decoder model for patients with greater than 150 trials (n = 15). Data Table corresponds to Figure 4b.

| Patient | Win Probability (%) | | | | | | | | | | |
| --- | --- | --- | --- | --- | --- | --- | --- | --- | --- | --- | --- |
|  | 0 | 0.1 | 0.2 | 0.3 | 0.4 | 0.5 | 0.6 | 0.7 | 0.8 | 0.9 | 1 |
| p01 | 50.0 | 81.8 | 66.7 | 82.4 | 71.4 | 80.0 | 60.0 | 71.4 | 85.7 | 60.0 | 40.0 |
| p02 | 75.0 | 64.6 | 73.7 | 50.0 | 79.2 | 72.7 | 81.8 | 84.6 | 82.4 | 90.9 | 72.7 |
| p03 | 43.8 | 79.2 | 77.9 | 85.7 | 60.0 | 80.0 | 60.0 | 66.7 | 57.1 | 73.7 | 40.0 |
| p04 | 66.7 | 79.2 | 72.7 | 66.7 | 54.6 | 90.0 | 70.0 | 73.7 | 66.7 | 60.0 | 18.2 |
| p06 | 60.0 | 70.0 | 75.0 | 68.8 | 79.2 | 72.7 | 81.8 | 84.6 | 82.4 | 90.9 | 72.7 |
| p07 | 40.0 | 79.2 | 73.7 | 72.7 | 71.4 | 66.7 | 82.4 | 71.4 | 66.7 | 70.8 | 33.3 |
| p08 | 63.6 | 79.2 | 71.4 | 76.3 | 73.7 | 73.7 | 78.3 | 54.6 | 66.7 | 50.0 | 50.0 |
| p09 | 30.0 | 64.6 | 66.7 | 77.9 | 66.7 | 70.8 | 78.3 | 76.3 | 61.1 | 63.6 | 63.6 |
| p10 | 54.6 | 57.1 | 71.4 | 81.8 | 76.3 | 63.6 | 71.4 | 66.7 | 63.6 | 82.4 | 33.3 |
| p13 | 30.0 | 57.1 | 76.3 | 61.1 | 63.6 | 66.7 | 81.8 | 71.4 | 73.7 | 70.0 | 33.3 |
| p14 | 57.1 | 73.7 | 71.4 | 66.7 | 43.8 | 79.2 | 77.9 | 71.4 | 71.4 | 81.8 | 50.0 |
| p15 | 66.7 | 77.9 | 71.4 | 71.4 | 66.7 | 79.2 | 72.7 | 73.7 | 54.6 | 66.7 | 33.3 |
| p16 | 33.3 | 81.8 | 57.1 | 73.7 | 50.0 | 90.0 | 50.0 | 43.8 | 66.7 | 54.6 | 18.2 |
| p18 | 73.7 | 79.2 | 77.9 | 85.7 | 30.0 | 57.1 | 76.3 | 71.4 | 85.7 | 77.9 | 75.0 |
| p19 | 71.4 | 76.3 | 66.7 | 54.6 | 90.0 | 79.2 | 71.4 | 71.4 | 66.7 | 57.1 | 66.7 |
| Mean | 54.4 | 73.4 | 71.3 | 71.7 | 65.1 | 74.8 | 72.9 | 70.2 | 70.1 | 70.0 | 46.7 |
| SD | 15.8 | 8.6 | 5.4 | 10.7 | 15.5 | 9.1 | 9.6 | 10.2 | 10.0 | 12.8 | 19.4 |

#### Extended Data Table 21: Decoder performance per patient, all trials

Performance (% accurately classified trials) for each decoder model for each patient. All trials are included.

| Patient | LV + DTW | LV + ED | PC + DTW | PC + ED | Best |
| --- | --- | --- | --- | --- | --- |
| p01 | 63.4% | 67.4% | 63.2% | 51.2% | 67.4% |
| p02 | 73.9% | 61.3% | 57.4% | 53.8% | 73.9% |
| p03 | 67.4% | 58.2% | 61.0% | 48.2% | 67.4% |
| p04 | 74.1% | 56.7% | 62.9% | 54.9% | 74.1% |
| p05 | 72.7% | 60.8% | 73.9% | 57.4% | 72.7% |
| p06 | 77.0% | 76.9% | 76.5% | 54.6% | 77.0% |
| p07 | 71.9% | 62.0% | 53.0% | 44.8% | 71.9% |
| p08 | 68.2% | 63.2% | 67.3% | 69.9% | 68.2% |
| p09 | 75.3% | 78.3% | 68.3% | 59.0% | 78.3% |
| p10 | 74.9% | 68.2% | 60.5% | 60.0% | 74.9% |
| p11 | 68.3% | 63.5% | 62.2% | 54.1% | 68.3% |
| p12 | 73.1% | 72.1% | 56.3% | 56.4% | 73.1% |
| p13 | 69.3% | 69.3% | 58.9% | 52.9% | 69.3% |
| p14 | 68.2% | 63.2% | 61.3% | 59.9% | 68.2% |
| p15 | 72.4% | 75.2% | 54.1% | 61.4% | 75.2% |
| p16 | 67.3% | 67.3% | 64.0% | 55.5% | 67.3% |
| p17 | 71.3% | 71.3% | 55.6% | 55.6% | 71.3% |
| p18 | 69.1% | 67.4% | 63.1% | 57.1% | 69.1% |
| p19 | 68.5% | 68.1% | 70.1% | 62.4% | 68.5% |
| p20 | 68.4% | 71.2% | 52.5% | 54.7% | 71.2% |
| Mean | 70.7% | 67.1% | 62.1% | 56.2% | 71.4% |
| SD | 3.34% | 5.85% | 6.46% | 5.19% | 3.34% |

#### Extended Data Table 22: Decoding results with and without beta

There was no difference between including or excluding β using LDS + DTW classifier model (p = 0.6, paired 2-tailed t-test) and results were highly consistent across patients (R^2^ = 0.75).

| Patient | β, γ, HFA | γ, HFA (no β) |
| --- | --- | --- |
| p01 | 62.4% | 63.4% |
| p02 | 71.4% | 73.9% |
| p03 | 67.4% | 67.4% |
| p04 | 74.1% | 74.1% |
| p05 | 74.7% | 72.7% |
| p06 | 72.3% | 77.0% |
| p07 | 71.9% | 71.9% |
| p08 | 68.2% | 68.2% |
| p09 | 71.9% | 75.3% |
| p10 | 74.9% | 74.9% |
| p11 | 68.3% | 68.3% |
| p12 | 74.3% | 73.1% |
| p13 | 69.3% | 69.3% |
| p14 | 71.2% | 68.2% |
| p15 | 72.4% | 72.4% |
| p16 | 67.3% | 67.3% |
| p17 | 71.3% | 71.3% |
| p18 | 69.1% | 69.1% |
| p19 | 69.5% | 68.5% |
| p20 | 68.7% | 68.4% |
| Mean | 70.5% | 70.7% |

#### Extended Data Figure 7: Decoder accuracy is robust to dropping regions/subcircuits

LV+DTW decoder results using a progressive strategy in which regions (A) or subcircuits (B) were progressively added to evaluate their contribution to accuracy (see Methods): (A) The average classification accuracy across subjects for a given number of regions when iteratively adding the region that improved classification accuracy the least. (B) The average classification accuracy across subjects for a all subsets of subcircuits.

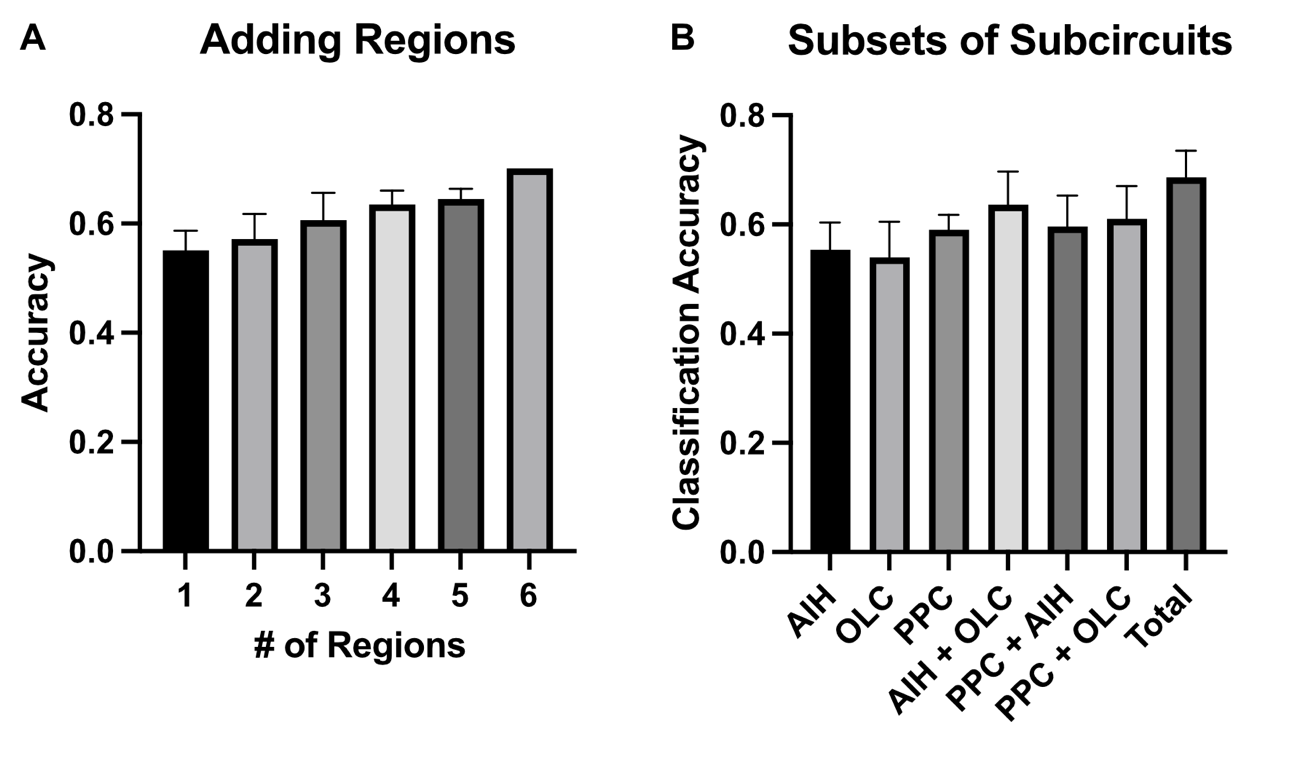

#### Extended Data Movie 1: Neural trajectories for gamble and safe bet choice

Movie showing the average latent variable trajectories (3 of 7 LVs) for safe bet and gamble trials for Patient 6, plotted over the 1 second prior to choice.

File: ED_Movie_1.mp4
